# Thermophilic Fungus Uses Anthraquinones to Modulate Ferrous Excretion, Sterol-Mediated Endocytosis and Iron Storage in Response to Cold Stress

**DOI:** 10.1101/2024.07.29.605589

**Authors:** Shuhong Li, Donglou Wang, Jiangbo He, Chunhua Liao, Zhangxin Zuo, Shenghong Li, Xuemei Niu

## Abstract

To date, there is no real physiological mechanisms for iron excretion in eukaryote, and no physiological “actuator” that can control all the three fundamental biologic processes of absorption, storage and excretion. Here we observed that the accumulation of anthraquinones by *Thermomyces dupontii* under cold stress can achieve this process. Through mutation analysis, we found that mutant Δ*An* deficiency in anthraquinones accumulated ferrous and total free iron due to adopting a rare lifestyle with no endocytosis but accumulation of membrane-derived vesicles. Anthraquinone complement indicated that the vesicles in Δ*An* could coat the extrinsic anthraquinone-induced granules to prevent contact with the fungal interiors. Detailed chemical investigation on Δ*An* led to characterization of a rare oxygen-free ergosterene with unstable nature in air as the major membrane steroid in Δ*An*, suggesting hypoxia inner in Δ*An* cells, consistent with dramatically low oxygen-consuming rates in Δ*An*. A series of physiological and metabolic analysis indicated anthraquinones were involved in exporting ferrous and promoting formation of oxygen-containing metabolites, including ergosterols for endocytosis and iron chelators for iron storage. Moreover, we found that both the anticancer agent mitoxantrone with well know-cardiotoxicity side effect and the major terpenoid-derived polycyclic aromatics from Danshen for treating cardiovascular disease showed potent ferrous transporting capabilities in human cancer cells. Our findings provide a novel insight into the underlying mechanisms of polycyclic aromatics in nature and pharmacology, and offer new strategy for developing potential therapeutics and agents for membrane transport, iron homestasis and anticold.

**Graphical Abstract:** Up to now, regulation of iron homeostasis by metabolites have rarely been characterized. Moreover, no excretory mechanisms for iron in fungi have been reported. In this study, we found that *Thermomyces dupontii* can accumulate a large amount of anthraquinones under cold stress. The anthraquinones can release free ferric ion, reduce ferric to ferrous ions, and export ferrous ions, greatly enhancing thermophilic fungus to survive in the bio-system environment. Furthermore, lack of the anthraquinones can inhibit oxygen-dependent ergosterol mediated endocytosis, leading to an self-imposed isolation via oxygen-free ergosterene-mediated cell membranes. Importantly, the well known anthraquinone compound Mitoxantraquinone for anticancer and the main terpenoid polycyclic aromatic metabolites in traditional Chinese medicine Danshen for the treatment of cardiovascular diseases both exhibit strong ferrous ions transport capabilities. Our findings provided new insights for developing potential therapies and drugs for iron homeostasis and drug delivery.

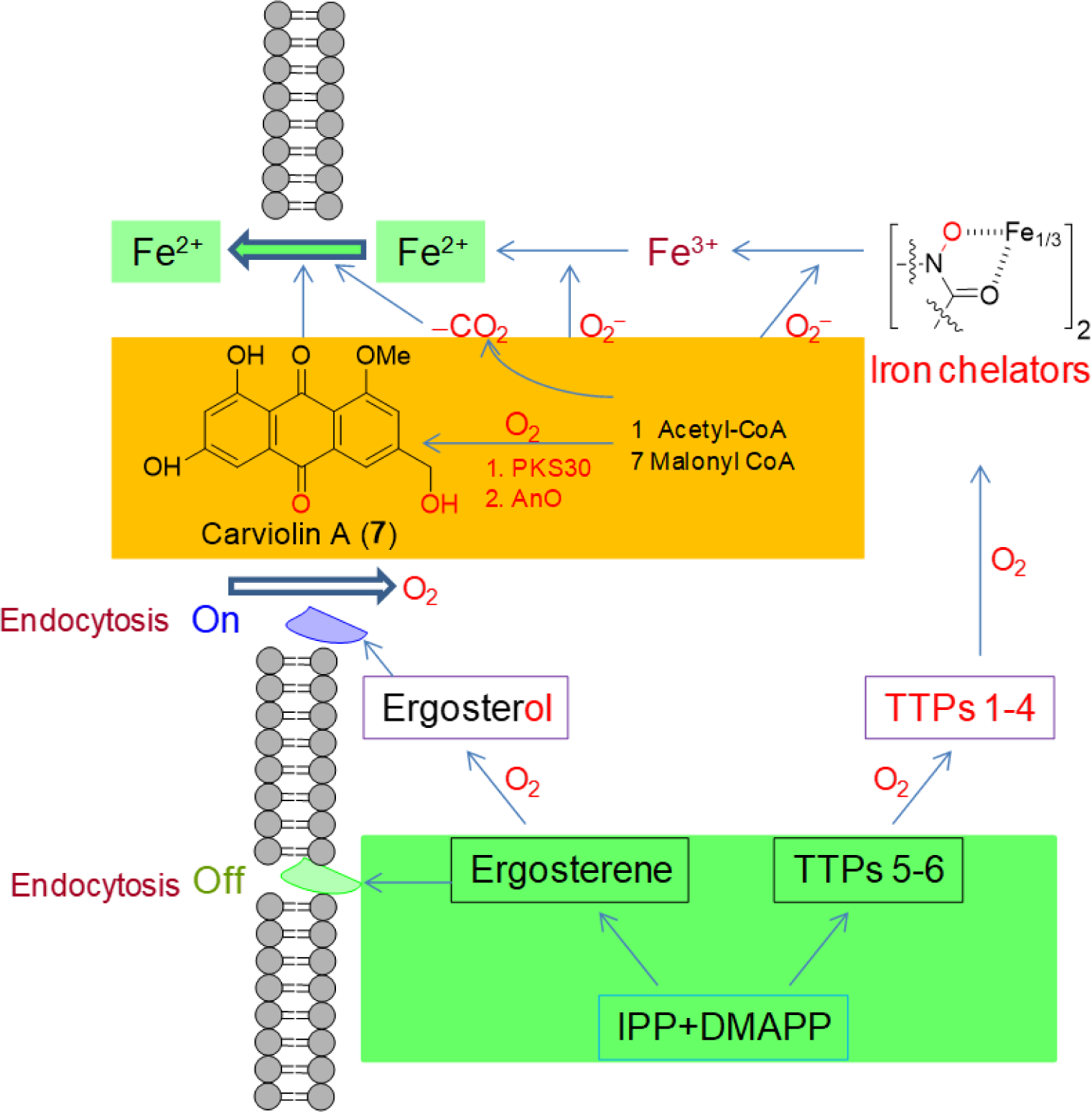

## Introduction

Anthraquinones are the largest group of quinoid natural pigments with rigid, planar, and aromatic chemical skeletons and are found in across various fungi, medicinal plants of various families, lichens and marine sources.^1−4^ Many anthraquinones exhibit yellow, orange, or brown pigmentation and are well known for their uses in the textile industry, paints, imaging photocleavable protecting groups, devices and biochips, foods, cosmetics, and pharmaceuticals.^4−7^ More prominently, the anthraquinone scaffold has been studied widely with respect to its pharmacological properties, such as its applications in antifungal, antiviral, antimalarial, antimicrobial, antiparasitic, anticancer, antiplatelet, antidiabetic, neuroprotective, laxative, and many more therapeutic settings. Anthraquinones inhibited topoisomerases, telomerase, protein kinases, matrix metalloproteinases (Zn and Ca-dependent neutral endopeptidases), and ecto-nucleotidases.^5−7^ In traditional medicines, anthraquinones have been used for many centuries, such as aloe emodin.^3^ However, the anthraquinones that are established to treat cancer, such as mitoxantrone, have been reported to display cardiotoxicity and other serious side effects.^5,7^ The underlying mechanism of such cardiotoxicity still remained largely unknown. Importantly, despite their widespread presence and biological activities, the natural functions of anthraquinones in native hosts have remained largely elusive.

Thermophilic fungi are unique eukaryotes capable of growing at high temperatures of 45−60 °C and are prevalent components of mycoflora in various composting systems.^8,9^ *Thermomyces*, a predominant thermophilic fungal genus, is phylogenetically closely related to common mesophilic fungi like *Aspergillus* spp. and *Penicillium* spp but contains largely reduced genomes. *Thermomyces* genus had only two species, *T. lanuginosus* and *T. dupontii* (*Penicillium dupontii*, and *Talaromyces thermophilus* before 2014).^8,9^ Both species exhibited optimal growth temperatures of 45−55 °C and a minimum growth temperature of 37 °C.^8^ Our recent studies have shown that both *Thermomyces* species displayed much darker colonies at minimum growth temperature of 37 °C compared to their optimal growth temperature of 45 °C.^10,11^ Among them, *T. lanuginosus* accumulated a melanin precursor, a polyketide derived dihydroxynaphthalene metabolite TDN.^10^ Through construction of the mutant Δ*Tdn* with the disruption of the polyketide synthase (PKS) gene *Tdn*, metabolic and transcriptional analysis, and a series of phenotypic and physiological bioassays, we found that TDN was involved in reinforcing fungal cell wall, inhibiting lipid formation and iron levels, and scavenging reactive oxygen species (ROS) to defend cold stress-induced ferroptosis.^10^ This study suggested that *Thermomyces* species should be an ideal model for deciphering physiological and biological functions of secondary metabolites in fungal adapt to temperature changes.

Interestingly, *T. dupontii* at the minimum growth temperature of 37 °C accumulated two types of metabolites, including nonribosomal peptides-terpenoid (NRP-TP) hybrid prenyl indole alkaloids (PIAs) and polyketide derived anthraquinones.^11^ Previous studies suggested that the major PIAs comprised of four complex PIAs, talathermophilins A−D (TTPs A−D, **1**−**4**), and two simple PIAs, TTPs E−F (**5**−**6**) (Fig. S1).^12–14^ Complex TTPs A−D are derived from simple TTPs E−F via both oxygenation and prenylation to form one more pyran ring fused with indole. All these PIAs can be oxygenated to form iron chelators for sequestering Fe^3+^, thus decreasing the levels of Fe^3+^ and total free iron.^14−19^ Unexpectedly, we observed significantly decreased levels of Fe^2+^ and total free iron levels in mutant Δ*TTP* without PIAs.^11^ Moreover, ROS levels and lipid formations were dramatically decreased, although superoxide levels were increased in the mutant Δ*TTP*.^11^ We postulated that the mutant Δ*TTP* might have used the anthraquinones to inhibit the levels of total free iron and ROS for protecting the mutant Δ*TTP* from cold-stress mediated ferroptosis.^10^ Therefore, in this study, we first evaluated anthraquinone levels in the mutant Δ*TTP* and characterized the major component among anthraquinones. Then, we investigated on natural functions of anthraquinones in *T. dupontii* under cold stress.

## EXPERIMENTAL PROCEDURES

### Mutant construction and culture conditions

*T. dupontii* was cultured in potato dextrose broths (PDB, potato (Kunming, China) 200 g L^-1^, glucose (Solarbio, Beijing, China) 10 g L^-1^) at 45 °C for 5 days. Mycelia were separated from broths via 2 layers of lens paper (10×15 cm, NEWSTAR Industry, Hangzhou, China) and collected for DNA extraction by using DNAiso Reagent (TaKaRa Biotechnology Co. Ltd, Dalian, China) as described in the manual. The concentration of the extracted DNA sample was determined using a Nanodrop system (NanoDrop, Madison, USA). A modified protoplast transformation method for genetic disruption of the *PKS* gene (Gene ID: Talth1_006479_t1) was applied using double-crossover recombination with the hygromycin-resistance gene (*hyg*) as a selection marker, followed by identification of desired mutants using diagnostic PCR.^20,21^ *Thermomyces dupontii* NRRL 2155 and mutants Δ*TTP* ^14^ and Δ*An* were cultured on potato dextrose agar (PDA, potato (Kunming, China) 200 g L^-1^, glucose (Solarbio, Beijing, China) 10 g L^-1^, agar (Solarbio, Beijing, China) 15 g L^-1^). All the bioassays were conducted in 9 cm diameter Petri dishes. Inocular of *T. dupontii* were cultured on PDA plates at 45 °C for 7 days or 37 °C for 14 days. PDA was used to determine the growth and other phenotypic traits of the fungus *T. dupontii* WT and different mutants. Plasmid pPK2.SUR.eGFP (a gift from Dr. Lianmin Liang, Yunnan University) was maintained in *Escherichia coli* strain DH5α (Takara, Shiga, Japan) and used to construct the recombinational plasmids.

### Ferric reduction assay

Carviolin A was isolated from *T. dupontii* WT according to our recent study.^10^ Carviolin A was dissolved in DMSO with the concentration 20 µM. 50 µL Carviolin solution was mixed with 50 µL 3-(2-pyridyl)-5,6-bis(5-sulfo-2-furyl)-1,2,4-triazin (#I291, Dojindo, Japan) in a total reaction mixture of 650 µL containing of 500 µL of 0.1 M acetate buffer (pH 4.4), and 50 µL of freshly prepared 1.0 mM FeCl_3_. Fe^3+^ reduction was assayed spectrophotometrically at 593 nm, monitoring the absorbance after 5, 30, 60 min of incubation with the iron reaction mixture. The real absorption values of the sample reactions were calculated differences between after and before joining 3-(2-pyridyl)-5,6-bis(5-sulfo-2-furyl)-1,2,4-triazin. Solutions without Carviolin A was used as a control. A standard curve was developed using ferrous sulfate. FeSO_4_ solutions were prepared at different concentrations of 0, 10, 20, 30, 40, 50, 60, 70, 80, 90, and 100 µM. A standard curve was developed in a total mixture containing 550 µL acetate buffer, 50 µL 3-(2-Pyridyl)-5,6-BIS (5-sulfo-2-furyl)-1,2,4-triazin, and 50 µL FeSO_4_. The calculated formula for the standard curve is y=0.0015x+0.0002, where y is the absorbance value and x is the Fe^2+^ concentration.^22^

### Metabolic analysis

Thermophilic fungus *T. dupontii* WT and two mutants Δ*TTP* and Δ*An* were cultured on PDA medium for 7 days, and the mycelia were inoculated into 500 mL flasks each containing 250 mL of PDB and incubated at 45 °C or 37 °C for 1−7 days on a rotary shaker (180 rev./min). The fermentation broths were exhaustively extracted overnight with 250 mL ethyl acetate (1:1 v/v) and the organic layers were concentrated to dryness under reduced pressure. The dried organic residue was dissolved in 1 mL methanol, filtered through 0.22 μM membranes, and analyzed by HPLC-MS performed on a Q Extractive Focus UPLC-MS (Thermo Fisher Scientific, USA) with a PDA detector and an Orbitrap mass detector (Shiseido, 5 μM, 4.6 mm x 250 mm, CAPCELL PAK C18 column) using positive and negative mode electrospray ionization. The total flow rate was 1 mL min^−1^; mobile phase A was 0.1% formic acid in water; and mobile phase B was 0.1% formic acid in acetonitrile. The column temperature was maintained at 40 °C. The injection volume for the extracts was 10 μL. The liquid chromatography (LC) conditions were manually optimized on the basis of separation patterns with the following gradient: 0–2 min, 10% B; 10 min, 25% B; 30 min, 50% B; 35 min, 90% B; 36 min, 95% B; 40 min, 95% B; 40.1 min, 10% B; and 45 min, 10% B. UV spectra were recorded at 196–400 nm.^10,11^ The data were analyzed and processed using Compound Discoverer 3.0 software.

### Iron measurement

Iron level (Fe^2+^ and Fe^3+^) was evaluated by Iron assay kit (#I291, Dojindo, Japan). The mycelia were separated from broths via 2 layers of lens paper (10×15 cm, NEWSTAR Industry, Hangzhou, China). In Brief, 100 mg mycelia was collected and washed with cold PBS three times. The fungal mycelia was resuspended in iron assay buffer and homogenized using the homogenizer sitting on ice. Then, the mixture was centrifuged at 12000 rpm for 10 min at 4 °C, and 100 μL supernatant was collected for detection. For extracellular iron level, 1 mL filtrate from the filtered broth were collected and centrifuged at 12000 rpm for 10 min at 4 °C. One hundred μL supernatant were collected for detection. Next iron assay buffer /reducer was added into the collected supernatant, mixed, and incubated according to the instruction. Finally, the incubated solution with iron probe for 1 h in the dark was immediately measured on microplate reader at OD = 593 nm.

### Fungal iron efflux bioassay

To further evaluate the iron efflux in *T. dupontii*, WT and Δ*An* were cultured on PDA medium for 7 days. Then the mycelia were inoculated into 250 mL flasks each containing 125 mL of PDB with 0.1 mM FeSO_4_ and incubated at 45 °C for 4 days on a rotary shaker (180 rev./min). One hundred mg of mycelia were used for iron level detection and another 100 mg were washed with sterilized PBS three times. The washed mycelia were inoculated in 100 mL modified Martin Medium (20 g/L glucose, 2 g/L NaNO_3_ (Nanjing Reagent, Nanjing, China), 0.5 g/L MgSO_4_•7H_2_O (Solarbio, Beijing, China), 1g/L KH_2_PO_4_ (Solarbio, Beijing, China)) and incubated at 37 °C (180 rev./min) for 2 days. Finally, the mycelia were separated from Martin medium broths via two layers of lens paper (10×15 cm, NEWSTAR Industry, Hangzhou, China). The filtered broths were used for the evaluation of iron levels.

### Congo Red bioassay

For acidic property test, the 9 mm fungal colonies of WT and Δ*An* were inoculated at 37 °C for 9 days on the plates of PDA medium supplemented with Congo red (0, 0.1, 0.2, 0.3 mg/mL). The diameter of each colony and acidic zone was measured. The experiments were performed at least three replicates.

### Conidial yield and spore germination

To compare the conidial yield and spore germination of the WT and Δ*An*, colonies of each strain initiated as mentioned above were incubated on PDA plates at 45 °C for 7 days and 37 °C for 12 days. The colonies were individually washed into 10 mL sterile water, which was then filtered through four layers of lens paper (10×15 cm, NEWSTAR Industry, Hangzhou, China) to remove hyphae. Then calculating the concentration of the conidial suspension by microscopic counting on a hemocytometer, 100 μL aliquots of conidial suspensions (1×10^8^ spores/mL) of the WT and the Δ*An* were incubated in 1 mL liquid YG medium containing 5 g/L yeast extract and 20 g/L glucosefor 8 h to 10 h at 45 °C or 37 °C to assay conidial germination rates. These experiments were performed in triplicate.

### Transcriptome analysis

Extraction of total and RNA from *T. dupontii* strains including mutant strains were extracted using the TRIzol® method following the manufacturer’s protocol (BGI-Shenzhen, China). The concentration of the extracted RNA samples was determined as described above, and the integrity of the RNA was examined by the RNA integrity number (RIN) using an Agilent 2100 bioanalyzer (Agilent, Santa Clara, USA).

RNA library was validating on the Agilent Technologies 2100 bioanalyzer for quality control. The double stranded PCR products above were heated denatured and circularized by the splint oligo sequence. The single strand circle DNA (ssCir DNA) was formatted as the final library. The final library was amplified with phi29 (Thermo Fisher Scientific, MA, USA) to make DNA nanoball (DNB) which had more than 300 copies of one molecular, DNBs were loaded into the patterned nanoarray and single end 50 bases reads were generated on BGISEQ500 platform (BGI-Shenzhen, China). Three biological replicates were analyzed for each sample. Low quality reads (Phred≤20) and adaptor sequences were filtered out, and the Q20, Q30 and total raw reads of clean date were calculated. Transcript abundances (FPKM) were provided by BGI after sequencing and differentially expressed genes were calculated using R software. All genes with P-value≤0.05 and log2 (fold_change) ≥1were considered significantly differentially expressed and used for KEGG pathway enrichment analysis.

### Assays for endocytosis and Fe^2+^ chemical probe in live cells

Sterilized cover-slips were half inserted into the PDA medium at a 45° angle slant and the 9 mm fungal colonies were inoculated onto PDA medium and cultivated at 45℃ for 3 days and at 37 ℃ for 5 days until the aerial hyphae grew onto and covered the cover-slips.To evaluate endocytosis, the cover-slips with aerial mycelia was incubated with FM4-64 staining solution (SynaptoRed C2, Biotium, Fremont, CA, USA; 10 μL of FM4-64 was diluted to a final concentration of 4 μM in 50 mM Tyrode solution) and immediately placed under a fluorescence microscope (Nikon, Tokyo, Japan) for observation.^23^ To evaluate Fe^2+^ in live cells, the aerial hyphae with cover-slips were incubated in 1 µM SiRhoNox-1(HY-D1533, MCE) for 1 h in the dark at room temperature prior to observations under fluorescence microscope.^24^

### Transmission Electron Microscopy (TEM)

For TEM analysis, inoculating WT and Δ*An* strains into PDB medium at 45 °C or 37 °C for 24 h. The mycelium shaken for 24 h was immediately placed in an EP tube containing 1 mL 2.5% glutaraldehyde for fixation. The samples were fixed overnight at 4 ℃ using 2.5% glutaraldehyde in 0.1 M PB (pH 7.2), then washed with 0.1 M PB (pH 7.2) three times for 7 min. Afterward, samples were postfixed with 1% KMnO4 for 4 h at 4 ℃, then washed with ddH_2_O three times for 7 min, followed by serial ethanol dehydration and acetone transition for 5 min, embedding in Epon 812 resin, polymerization at 60 ℃ for 48 hours. Serial sections of uniform thicknesse, 800 nm for semithin sections and 60 nm for ultrathin sections, were made using a leica EM UC7 ultramicrotome. Ultrathin sections were then loaded onto Cu grids and double stained with 2% uranyl acetate and lead citrate before observations employing a JEM-1400 Plus transmission electron microscope at 80 kv.

### Isolation and characterization of ergosterene from mutant Δ*An*

Thirty L of PDB fermentation broths of the Δ*An* mutant were subjected to vacuum concentration to obtain 3 L and extracted with ethyl acetate. The ethyl acetate portion was evaporated and dried to generate residue, which was dissolved in a small amount of methanol. The crude extract of the mutant was subjected to column chromatography using silica gel 60 (Merck, 200−300 mesh) and eluted with a gradient of 60−100% H_2_O−MeOH (60−100%) using reverse phase (RP18) flash chromatography. Subsequently, the sample was further purified using column chromatography on Sephadex LH-20 with acetone. Finally, ergosterene (2.3 mg) was obtained by purification with silica gel 60 eluting with Petroleum ether/dichloromethane (3:1). NMR experiments were carried out on a DRX-500 spectrometer with TMS as internal standard.

### Chemical complementation

Fungal strains was cultured on PDA for 7 days, and then 1×10^7^ fungal spores were washed down and inoculated into each 250 mL flask containing 125 mL of PDB. Tanshinone IIA (TS IIA), tanshinone I (TS I) and cryptotanshinone (CTS) were purchased from Chengdu University of Traditional Chinese Medicine. For chemical complementation, carviolin A, tanshinone IIA (TS IIA), tanshinone I (TS I) or cryptotanshinone (CTS), dissolved in DMSO was added into fermentation broth after inoculation for 24 hours at a final concentration of 20 μM.

### Human cancer cell bioassay

Human hepatocellular carcinoma HepG2 cells (HepG2) were provided by Yunnan Suli Biopharmaceutical Company. HepG2 cells were cultured in DMEM (Gibco, Carlsbad, CA, USA) supplemented with 10% fetal bovine serum (BI, Kibbutz Beit Haemek, Israel) and 1% penicillin/streptomycin (Gibco); HepG2 cells were cultured at 37 ℃ in a 5% CO_2_ humidified environment. About 1×10^6^ HepG2 cells in 5 mL of DMEM were added to 25 cm^2^ dish and carviolin A (CA), Tanshinone IIA (TS IIA), Tanshinone I (TS I), cryptotanshinone (CTS) or mitoxantrone (MITO) dissolved in DMSO was added into dulbecco’s modified eagle medium (DMEM) at a final concentration of 5 μM and then incubated 48 h at 37 °C in 5% CO_2_. Solvent DMSO was used as a control. Then HepG2 cells were collected and washed with cold PBS three times and the cell viability was evaluated under a microscope with a cell counter plate. The iron levels (Fe^2+^ and Fe^3+^) in cells were evaluated with iron assay kit (#I291, Dojindo, Japan) according to the above method.

### ROS/superoxide assay

ROS/Superoxide Detection Assay Kit (#ab139476, Abcam, Britain) was used to directly monitor of reactive oxygen species (ROS) and Superoxide in mycelia. Twenty mg mycelia was harvested from PDA in 2 mL centrifuge tube and were treated with 500 μL of ROS/Superoxide Detection Solution and incubated for 60 min at 37 ℃ or 45 ℃ in the dark. The stained mycelia were analyzed by microplate reader. Oxidative Stress Detection Reagent (Green, Ex/Em 490/525 nm) was used for evaluation of total ROS, and Superoxide Detection Reagent (Orange, Ex/Em 550/620 nm) for Superoxide.

### Oxygen Consumption Rate Test

Mitochondrial Stress Test Complete Assay Kit (# ab197243, Abcam, Britain) was applied to detect oxygen consume rates (OCR) according to the manufacturer’s instructions. About 10 mg fungal hyphae were added into 1.5 ml centrifuge tube with 1 mL PDB medium and mixed in vortex oscillator. Then, 100 μL was inoculated into 96-well microplate (1 mg/Well). Subsequently, 8 μL of reconstituted extracellular O_2_ consumption reagent and 100 µL of pre-warmed high sensitivity mineral oil (37 °C) were added to each well. Plates were read at 37 °C in kinetic mode on the CLARIOstar energy metabolism monitor for 4 hours (read every 2 minutes, Ex/Em = 380/650 nm). The basal OCR levels were calculated according to the slope of the kinetic curve.

### Bioinformatic analysis

Using the protein sequences of PKS30 in *T. dupontii* as queries, we performed a reciprocal blast against gene catalog protein in MycoCosm database using the blastp algorithm. Only the alignment hit coverage (%) and sequence identity (%) of a sequence greater than 40 was retained, and the corresponding fungi species was considered to have homologous *PKS30* gene. Analogues of *PKS30* in 636 fungal genomes from 2165 fungal genomes belonging to 134 fungal genera 749 fungal genera were finally retrieved. The distribution of PKS30-containning fungal genomes were mapped to the genus-levels fungal phylogenetic tree.

### Data analysis

Data from three biological repeated experiments were expressed as means ± SD, which were analyzed by one-way analysis of variance followed by Tukey’s multiple comparison test, with p values < 0.05 (*), p values < 0.01 (**), p values < 0.01 (***), considered statistically significant. All statistical analyses were conducted using GraphPad Prism ver. 8.00 for Windows (GraphPad Software, San Diego, CA, USA).

## RESULTS

### Mutant Δ*TTP* at 37 °C accumulated anthraquinones in mycelia and displayed elevated levels of Fe^2+^ and total free iron in broths

Mutant Δ*TTP* at 37 °C displayed a dark-colored pigment similar to WT at 37 °C (Fig. 1A). Metabolic analysis indicated that the mutant Δ*TTP* at 37 °C showed an extra HPLC peak at retention time of 12.6 min not present at 45 °C, also similar to WT (Fig. 1A). This peak was characterized to be carviolin A (**7**), an anthraquinone metabolite, according to its quasi-molecular ion peak at *m/z* 299.0557 [M−H]^−^ for a molecular formula of C_16_H_12_O_6_ in the negative HRESI mass spectrum, and the UV absorptions (Figs. S2−S3).^25^ Further chemical investigation on a 30 L fermentation cultures of the mutant Δ*TTP* led to isolation of metabolite **7**. The ^1^H and ^13^C NMR data of **7** finally confirmed the structure of **7** as carviolin A (Figs. S4−S5).

**Figure 1.**
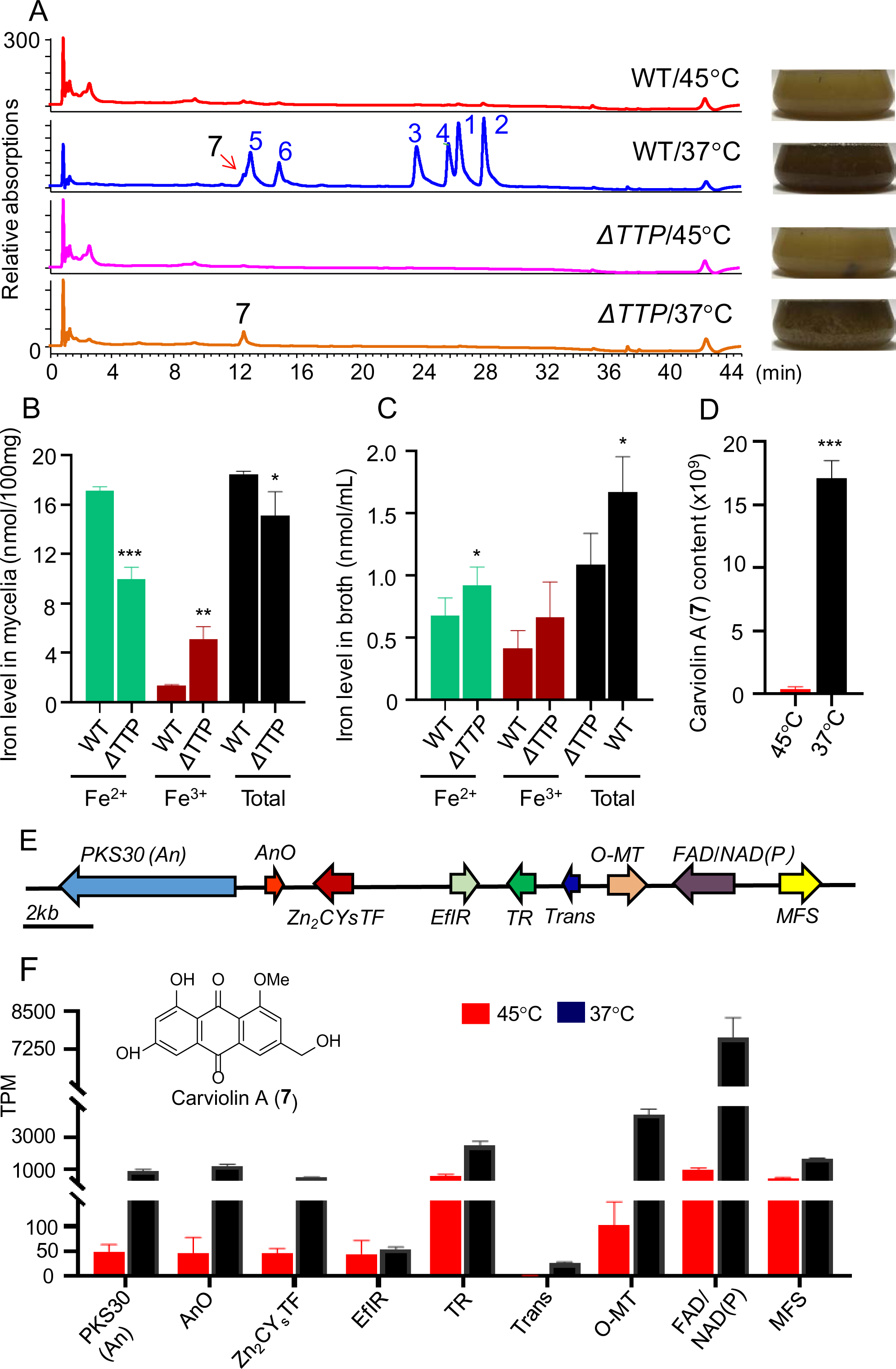
Comparison of phenotypes, metabolic profiles and iron levels of WT and the mutant Δ*TTP* and transcriptional analysis of the gene cluster responsible for anthraquinone biosynthesis at fungal minimum growth temperature of 37 °C and optimal growth temperature of 45 °C. A) Phenotype analysis visually illustrates the presence of dark colored pigments in both mutant Δ*TTP* and WT at minimum growth temperature of 37 °C, a distinct contrast to their respective strains at optimal growth temperature of 45 °C, and metabolic analysis by HPLC-DAD indicates that while WT exhibits 7 extra metabolites at 37 °C compared to 45 °C, the mutant Δ*TTP* (highlighted in red) produces an additional metabolite (**7**) at 37 °C compared to 45 °C. B−C) Comparison of iron levels in mycelia (B) and broths (C) between WT and the Δ*An* mutant at 37 °C. D) Quantitative analysis focuses on metabolite **7**, known as carviolin A, in the fungal WT at 37 °C vs. 45 °C. E) The organization of the *An* gene cluster responsible for the biosynthesis of carviolin A (**7**). Key genes involved include Anthrone oxygenase gene (*AnO*), O-methyltransferase gene (*O-MT*), FAD/NAD(P)-binding domain-containing protein gene (*FAD/NAD(P)*), Zn_2_Cys_6_ transcription factor gene (*Zn_2_CysTF*), ungal_trans domain-containing protein gene (*Trans*), thioredoxin-like domain-containing protein gene (*TR*), and MFS transporter gene (*MFS*). F) Transcriptional analysis of the genes in the *An* gene cluster in the fungal WT at 37 °C vs. 45 °C shows that temperature reduction induces gene expression related to anthraquinone biosynthesis. p < 0.05 (*), p < 0.01 (**), p < 0.01 (***).

The contents of carviolin A (**7**) in Δ*TTP* was 2.08-fold higher than those in WT at 37 °C (Fig. S6), consistent with that the content of anthraquinone carviolin A (7) was positively correlated to the dark color of the WT fermentation broth. Detailed metabolic analysis of Δ*TTP* and WT revealed that another nine anthraquinones, including four anthraquinone dimmers, were remarkably elevated in Δ*TTP* vs. WT at 37°C (Fig. S7). The increase in anthraquinone contents in Δ*TTP* might explain the decreased lipid levels in Δ*TTP* compared to WT due to the fact that lipid formation and anthraquinone biosynthesis share the same precursors, acetyl-CoA and malonyl-CoA.^1-3^ Previous studies have reported that anthraquinone metabolites exhibit strong ROS scavenging activities while enhancing superoxide formation.^26,27^ Notably, superoxide can convert Fe^3+^ to Fe^2+^, and Fe^2+^ is oxidized to Fe^3+^ upon the conversion of hydrogen peroxide to the highly ROS hydroxyl radical in a process known as the Fenton reaction.^27^ In this study, we also found that carviolin A (**7**) could reduce Fe^3+^ to Fe^2+^ (Fig. S8). This raised a question: Either the Fe^2+^ levels or the ROS levels should have increased in Δ*TTP* at 37 °C, why were both decreased?

Since anthraquinone pigments were excreted from mycelia into fungal broths, we deduced that anthraquinone excretion might affect free iron levels in and out of the Δ*TTP* mutant. We compared the levels of free iron ions, including Fe^2+^, Fe^3+^ and total free iron, in mycelia and broths between WT and Δ*TTP* at 37 °C. As expected, in mycelia, the mutant Δ*TTP* exhibited significantly decreased levels of Fe^2+^ and total free iron compared to WT (Fig. 1B), consistent with the previous study.^11^ Interestingly, in broth of the mutant Δ*TTP* displayed dramatically increased levels of Fe^2+^ and total free iron, compared to WT (Fig. 1C), suggesting that anthraquinones in Δ*TTP* should export Fe^2+^ from mycelia to broths.

### Loss of anthraquinones elevated ROS levels and decreased conidial formation in mutant Δ*An* at 37°C

Upon examination of metabolic and transcriptional profiles in the WT at 37 °C vs. 45 °C, the contents of carviolin A (**7**) in WT at 37 °C was 44.02-fold higher than that at 45 °C (Fig. 1D) and one *PKS* gene *An* (*PKS30*) putatively responsible for anthraquinone biosynthesis was strongly up-regulated at 37 °C with 16.4 times higher than that at 45 °C. Furthermore, several other biosynthetic genes within the *An* gene cluster, encoding Anthrone oxygenase (AnO), O-methyltransferase (O-MT), and FAD/NAD(P)-binding domain-containing protein (FAD/NAD(P)), were also significantly upregulated at 37 °C, with 7.3- 42.7 times higher than those at 45 °C (Figs. 1E−1F, Fig. S9). These findings strongly suggested that the *An* gene cluster in *T. dupontii* played a pivotal role in the fungal response to temperature reduction from 45 °C to 37 °C. Moreover, genes embedded within the *An* cluster, such as Zn_2_Cys_6_ transcription factor gene (*Zn_2_CysTF*), fungal_trans domain-containing protein gene (*Trans*), thioredoxin-like domain-containing protein gene (*TR*), and MFS transporter gene (*MFS*), were also significantly upregulated in WT at 37 °C vs. 45 °C, further supporting the up-regulation of *An* gene cluster in response to temperature reduction.

To characterize the role of anthraquinones in the fungus during temperature reduction, we successfully generated a mutant with the deletion of gene *An* (Δ*An*) using a modified homologous recombination system for *T. dupontii* (Fig. S10 and Table S1).^20,21^ Phenotypic analysis revealed the disappearance of the dark colored pigment in Δ*An* compared to WT at 37 °C (Fig. 2A). Metabolic analysis further confirmed loss of the target anthraquinone metabolite, carviolin A (**7**), and its derived metabolites in Δ*An* at 37 °C (Fig. 2B, Fig. S11). Meanwhile, Δ*An* exhibited a notably decreased superoxide levels but elevated ROS levels compared to WT (Figs. S12A−B), aligning with the notion that anthraquinones can increase superoxide levels but attenuate ROS levels in Δ*TTP*. Importantly, mutant Δ*An* at 37 °C showed strongly decreased conidial formation by 89.93% and spore germination rates by 25.33% compared with WT at 37 °C (Figs. S12C−D).

**Figure 2.**
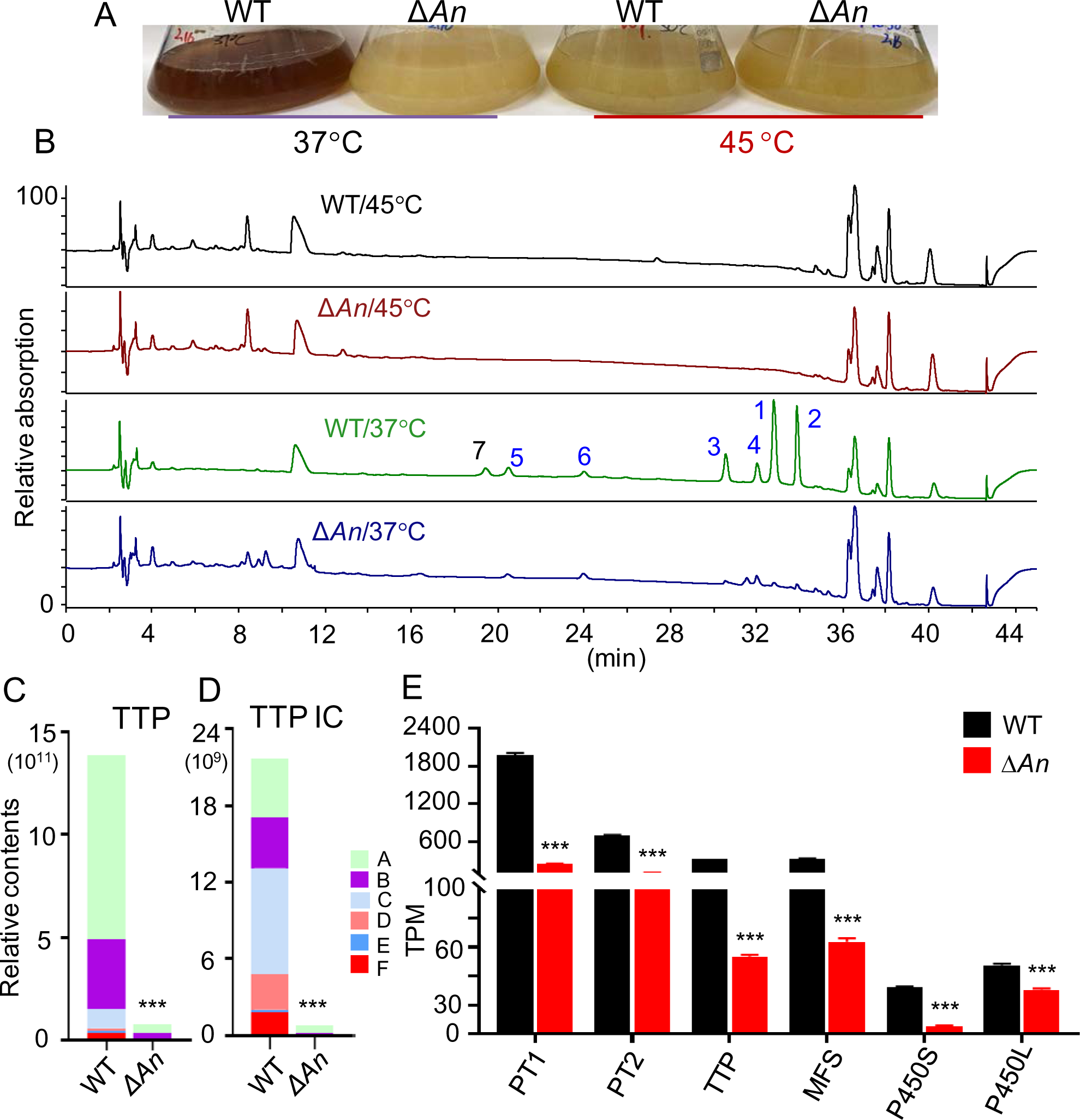
Phenotypic, metabolic and transcriptional analysis of the mutant Δ*An* and WT. A−B) Comparison of the broth pigments (A) and metabolic profiles (B) between WT and the mutant Δ*An* at 37 °C vs. 45 °C. C−D) Comparison of TTP levels (C) and their derived iron chelators (TTP IC, D) between WT and the mutant Δ*An* at 37 °C vs. 45 °C. E) Transcriptional analysis of all biosynthetic genes for TTP biosynthesis between WT and mutant Δ*An* at 37 °C. p < 0.05 (*), p < 0.01 (**), p < 0.01 (***).

### Mutant Δ*An* mostly lost TTPs and derived iron chelators

Surprisingly, we observed dramatically decreased TTPs contents in metabolic profile of the Δ*An* mutant, compared to WT at day 7 (Fig. 2C). Detailed comparisons of the contents of TTPs and their derived iron chelators between Δ*An* and WT at 37 °C for 7 consecutive days were conducted (Fig. S13). Notably, complex PIAs, oxygen-containing TTPs A−D (**1**−**4**), consistently showed a substantial reduction (ranging from 7.3−42.7 fold less) in Δ*An* compared to WT. The simple PIAs, TTPs E−F (**5**−**6**) also exhibited significant decreases in Δ*An* compared to WT at most of the time points. Importantly, all the iron chelators derived from TTPs A−F (**1**−**6**) were drastically reduced in Δ*An* (Fig. 2D). We further conducted transcriptional analysis of Δ*An* and WT at 37 °C. Genes *AnO*, *O-MT*, and *TTP* were among the top ten down-regulated genes in the Δ*An* mutant compared to WT at 37 °C (Fig. S14). All the genes in *TTP* gene cluster, including two prenyltransferase genes *PT1* and *PT2*, two *P450* genes and one transport gene, were strongly down-regulated in Δ*An* vs. WT (Fig. 2E), consistent with the dramatically decreased levels of TTPs in Δ*An*. These results suggested that loss of anthraquinones could decrease the formation of TTPs and derived iron chelators in the Δ*An* mutant. Thus, the key difference between mutants Δ*An* and Δ*TTP* is that one lacks anthraquinones and the other produces a large amount of anthraquinones.

### Anthraquinones induced Fe^2+^ efflux and increased Fe^2+^/Fe^3+^ ratio in mutant Δ*An*

Iron levels, including Fe^2+^, Fe^3+^ and total free iron, were evaluated in the mycelia and broths of Δ*An* and WT at 37 °C (Figs. 3A−3F). Notably, in mycelia, the Δ*An* mutant displayed significantly higher levels of Fe^2+^, Fe^3+^ and total free iron compared to WT. Remarkably, in broths, the Δ*An* mutant exhibited drastically lower levels of Fe^2+^ and total free iron than WT at 37 °C (Fig. 3B). The Fe^3+^ levels of the broths in Δ*An* was a little lower compared to WT. It was unreasonable to attribute the lower levels of Fe^2+^ and total free iron to iron uptake since Δ*An* was unable to compete with WT due to significant reduction in TTPs and deficiency in anthraquinone biosynthesis. The remarkably high levels of Fe^2+^ and total iron in the broths of WT must be ascribed to Fe^2+^ efflux. To further verify the conclusion, Fe^2+^ efflux bioassays for WT and Δ*An* were conducted with the modified Martin medium without strains as blank control. As shown in Fig. S15, both WT and the mutant strain Δ*An* displayed significantly increased levels of Fe^2+^ and total free irons in the broths compared to the blank medium without strains, indicating that the fungal strains could elevate the iron contents in the environments. Moreover, WT exhibited more Fe^2+^ and total free irons than the mutant Δ*An*, suggesting that loss of anthraquinone biosynthesis specifically inhibited Fe^2+^ efflux from fungal mycelia to the broths.

**Figure 3.**
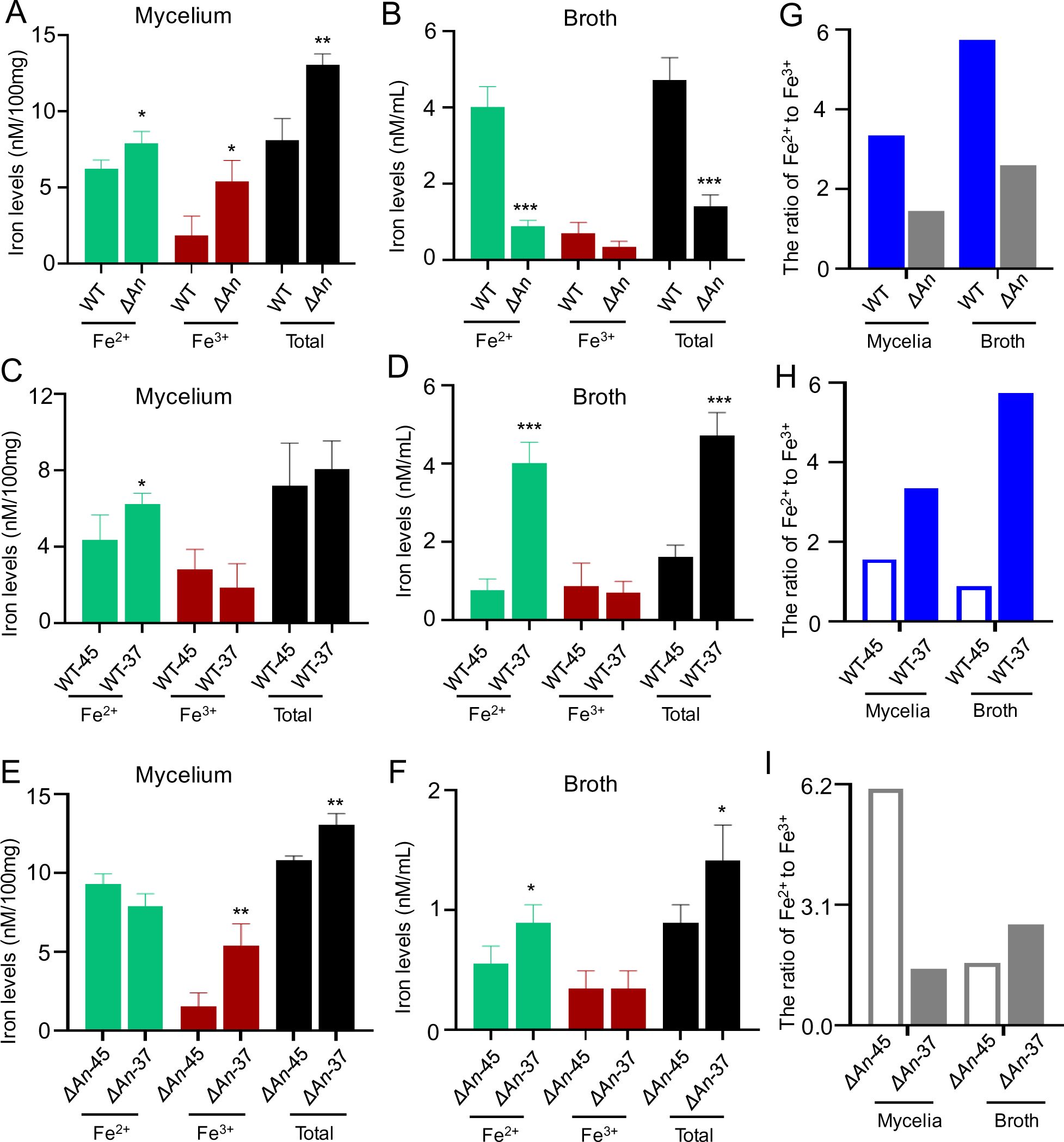
Evaluation of iron levels in mycelia and broths of WT and the mutant Δ*An* at 37 °C and at 45 °C indicated the role of anthraquinones in improving Fe^2+^ efflux and Fe^2+^/Fe^3+^ ratios in fungal response to cold stress. A−B) Comparison of iron levels in mycelia (A) and broths (B) between WT and the Δ*An* mutant at 37 °C. C−D) Comparison of iron levels in mycelia (C) and broths (D) of WT at 37 °C and 45 °C. E−F) Comparison of iron levels in mycelia (E) and broths (F) of the mutant Δ*An* at 37 °C and 45 °C. G) Comparison of the Fe^2+^/Fe^3+^ ratio in mycelia and broths between WT and the Δ*An* mutant at 37 °C. H) Comparison of the Fe^2+^/Fe^3+^ ratio in mycelia and broths of WT at 37 °C and 45 °C. I) Comparison of the Fe^2+^/Fe^3+^ ratio in mycelia and broths of Δ*An* at 37 °C and 45 °C. p < 0.05 (*), p < 0.01 (**), p < 0.01 (***).

However, it was a paradox that in mycelia, the mutant Δ*An* displayed higher levels of Fe^2+^ and total iron levels than WT at 37 °C, yet the Fe^2+^ efflux in the mutant Δ*An* was remarkably inhibited. These results suggested that another factor might be involved in the induction of Fe^2+^ efflux in the fungus at 37 °C. Considering that Δ*An* also displayed higher Fe^3+^ levels than WT at 37 °C, we evaluated the ratio of Fe^2+^/Fe^3+^ in both WT and Δ*An* at both temperatures (Fig. 3G). Notably, Δ*An* exhibited significantly lower ratio of Fe^2+^/Fe^3+^ than WT at 37°C in mycelia and broths. Moreover, WT exhibited significantly higher ratio of Fe^2+^/Fe^3+^ at 37 °C than 45°C in both mycelia and broth (Fig. 3H), indicating that the fungus increased ratio of Fe^2+^/Fe^3+^ to response to cold stress. Mutant Δ*An* displayed strongly decreased ratio of Fe^2+^/Fe^3+^ in mycelia at 37 °C vs. at 45 °C (Fig. 3I), consistent with the above result that anthraquinones could reduce Fe^3+^ and increase the ratios of Fe^2+^/Fe^3+^.

Previous study reported that oxidation of Fe^2+^ by molecular oxygen under acidic conditions was kinetically slow.^28^ Because Congo red has been widely used as an acid indicator, Congo Red bioassay for evaluation of acidic conditions of the fungal strains were carried out. Our results showed that both WT and Δ*An* displayed acidic cycles (Fig. S16). In particular, WT exhibited larger acidic cycles than Δ*An*. All the results indicated that even in the presence of oxygen, the acidic conditions of fungal growth were in favor of Fe^2+^ instead of Fe^3+^, thus consistent with the fact that Fe^2+^ levels were higher than Fe^3+^ levels in the fungus.

### Anthraquinones improved intracellar dark granule formation and endocytosis

To investigate what was occurred in the Δ*An* mutant, where Fe^2+^ efflux was inhibited, we performed transmission electron microscopy (TEM) analysis, using WT as control. We observed that WT at 37 °C harbored more conspicuous black granules, which were significantly reduced in WT at 45 °C and in Δ*An* at both 37 °C and 45 °C (Figs. 4a−d, Fig. S17). In the periphery of WT mycelia, tiny morphological structures extruded from the WT cell walls were clearly observed. These extruded structures were significantly increased in the periphery of WT at 37 °C compared to 45 °C (Figs. 4a−b), aligning with the significant increase in Fe^2+^ efflux in WT due to temperature reduction. Noticeably, these extruded structures were drastically decreased in the periphery of the mutant Δ*An*, consistent with the remarkable decrease in Fe^2+^ efflux in Δ*An* compared to WT. Another compelling observation was the morphological change in mycelial features between WT and Δ*An*, where several inner membrane-derived vesicles appeared in all the Δ*An* mutant while not in WT (Figs. 4c−d).

**Figure 4.**
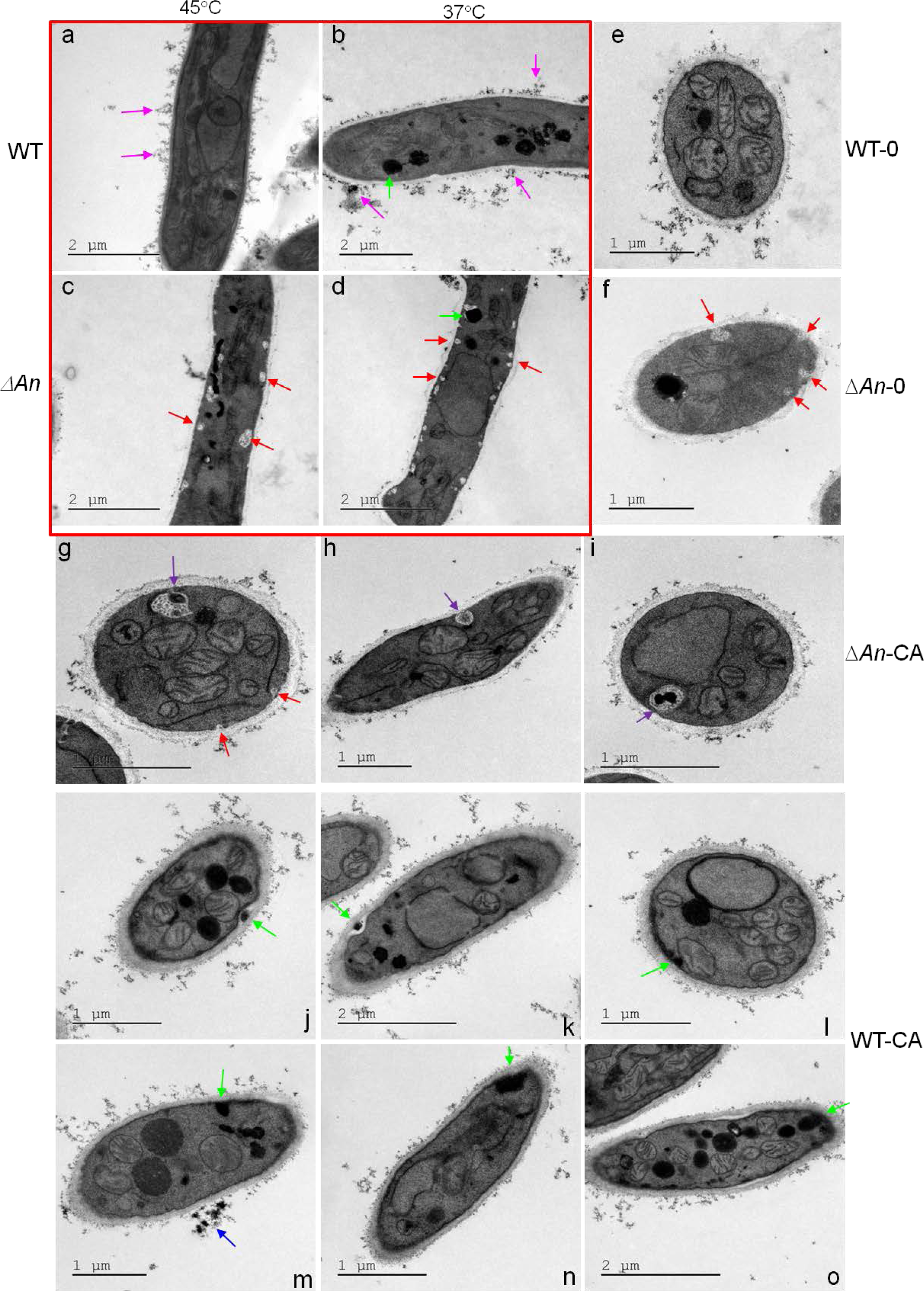
TEM analysis of the effects of anthraquinones on granule formation and fungal endocytosis. a−d) Comparison of WT and the mutant Δ*An* at 37 °C and 45 °C. Purple arrow: extruded material from WT; Green arrow: black granules; Red arrow: vesicles. e−f). Comparison of WT and Δ*An* treated with solvent DMSO. 0: DMSO. g−i) TEM analysis of Δ*An* treated with carviolin A (CA). Red arrow: white vesicle observed in Δ*An* treated with solvent; Purple arrow: a vesicle containing a black granule in Δ*An* treated with carviolin A. j−o) TEM analysis of WT treated with carviolin A. Green arrow: black endocytic granules observed externally and internally in WT treated with CA. Blue arrow: black granules formed in WT broth treated with carviolin A.

A chemical complementation of Δ*An* with 20 μM carviolin A (**7**) at 37°C was conducted with solvent as control. Blank solvent was used as control, and WT strain as control strain. TEM analysis showed that carviolin A treatment induced the appearance of black granules in both the Δ*An* mutant and WT (Figs. 4e−o), suggested that carviolin A was involved in the formation of black granules. In particular, WT under carviolin A treatment accumulated far more black granules than the Δ*An* mutant (Fig. S18). Intriguingly, the black granules were captured in a large vesicle in the Δ*An* mutant (13/18) but not in WT (0/18) (Figs. 4g−i), suggesting that the vesicles in the Δ*An* mutant functioned as quarantine organelles to prevent from external intrusion and maintain internal homeostasis. Importantly, WT treated with carviolin A displayed these black granules in various stages of endocytosis. Among them, membrane invagination was induced by the black granules in WT treated with carviolin A (4/18), while not with solvent (0/18). This observation suggested that exogenous carviolin A enhanced membrane invagination for influx. Conversely, endogenous carviolin A could promote membrane evagination for efflux.

To further evaluate the effect of anthraquinones on endocytosis, we stained both the Δ*An* mutant and WT with lipophilic styryl dye FM4−64, an endocytic tracer.^23^ In WT, FM4-64 gradually internalized and stained endosomes after 3−4 minutes of incubation at 37 °C and 45 °C, respectively (Fig. 5A). In contrast, only the plasma membrane next to hyphal wall was stained with FM4-64 in Δ*An* (Fig. 5A). Additionally, Δ*An* displayed much fewer dyed granules inside the hyphae than WT, suggesting that lack of anthraquinones could inhibit endocytosis. Further bioassay for oxygen consumption rates (OCR) indicated that the Δ*An* mutant displayed much lower OCRs than WT at 37 °C (Fig. 5B), consistent with the decreased endocytosis in Δ*An*. Chemical probe RhoNox-1 has been commonly applied for the evaluation of Fe^2+^ in live cells.^24^ We also carried out the Fe^2+^ assay in the live mycelia using RhoNox-1 according to the previous reference.^24^ We found that the chemical probe could enter WT strain and produce orange fluorescence, but failed in the mutant strain Δ*An* (Fig S19), consistent with the fact that the endocytosis was significantly inhibited in Δ*An*.

**Figure 5.**
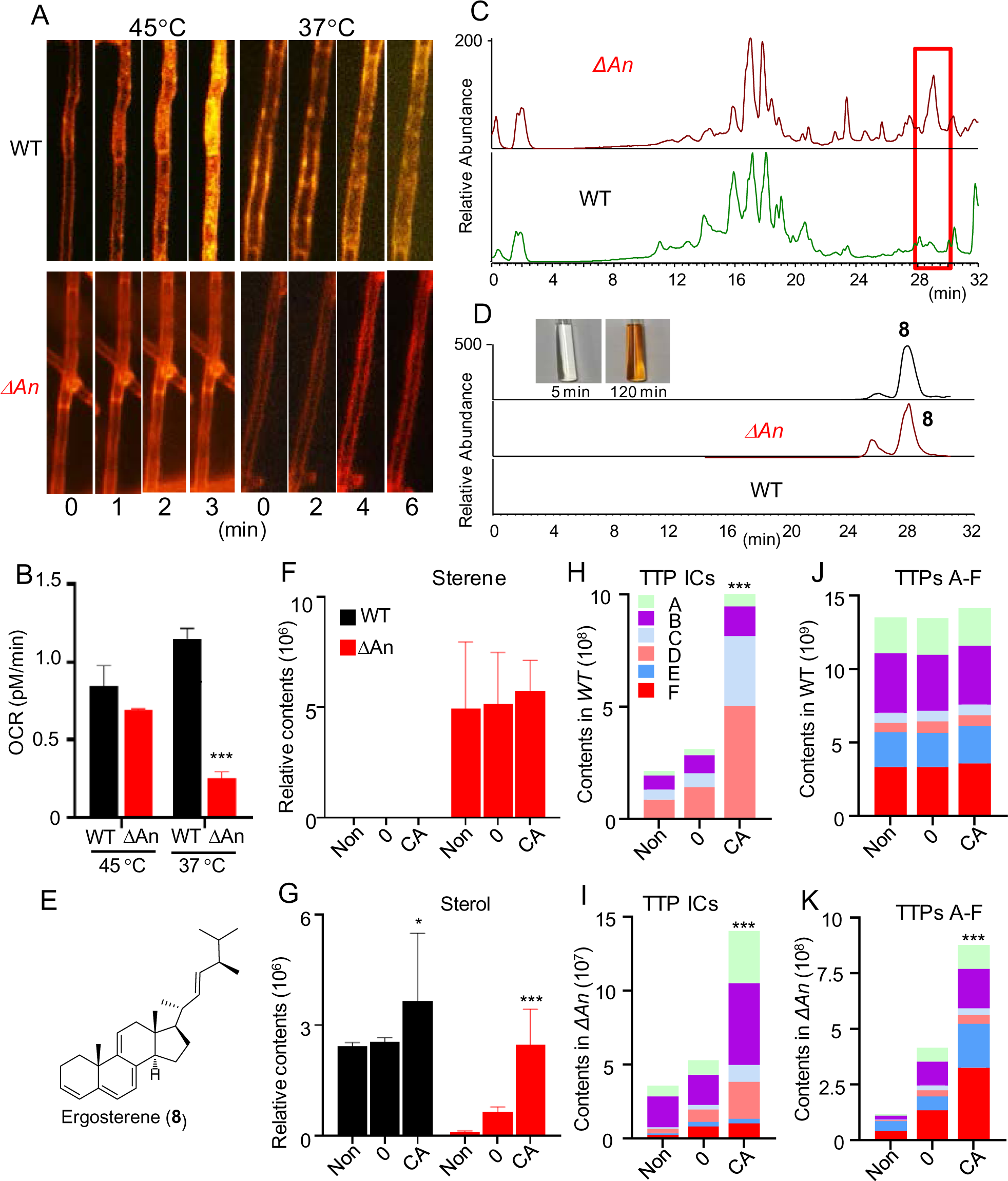
Evaluation the impact of anthraquinones on fungal endocytosis, oxygen consumption rates, membrane components, and iron chelators. A) Comparative analysis of endocytosis between WT and the Δ*An* mutant at 37 °C and 45 °C using the lipophilic styryl dye FM-464. B) Comparison of oxygen-consuming rates (OCRs) between WT and the mutant Δ*An* at both temperatures 37 °C and 45 °C. C) Metabolic profiles of WT and the Δ*An* mutant at 37 °C. D) HPLC analysis of ergosterene (**8**) in WT and the mutant Δ*An* at 37 °C. E−G) Comparison of the levels of ergosterene (**8**) and ergosterol in WT and the Δ*An* mutant treated with and without carviolin A at 37 °C. H−K) The effects of carviolin A on the levels of iron chelators (H and I) derived from TTPs A−F (J and K) in WT and the Δ*An* mutant treated with 20 μM carviolin A at 37 °C. p < 0.05 (*), p < 0.01 (**), p < 0.01 (***).

### An oxygen free ergosterene was characterized in mutant Δ*An*

To understand how membrane transport was hindered, we conducted comparative transcriptional and metabolic analyses. Transcriptional profiles revealed that polyketide biosynthesis enzymes, membrane transport, and ABC transporters were the top three down-regulated pathways in Δ*An* vs. WT (Figure S20A). This data aligned with the absence of anthraquinone biosynthesis and significant reduction in Fe^2+^ efflux, vesicle membrane transport and endocytosis. We also observed upregulation in steroid biosynthesis as well as metabolism of terpenoids and polyketides, both related to fungal membrane in Δ*An* vs. WT (Figure S20B). Subsequent metabolic profiling identified several low polar components, potentially assigned as steroids, which accumulated significantly in Δ*An* (Fig. 5C). To characterize the target metabolites in the Δ*An* mutant, chemical investigation was conducted on 30 L of the culture filtrate of Δ*An*. One major target metabolite (**8**) was eventually isolated. However, this metabolite (**8**) proved to be rather unstable and rapidly oxidized in the air, causing its color to change from colorless to brown within only one hour (Fig. 5D). Eventually, we managed to obtain the MS and 1D and 2D NMR data of this metabolite (**8**) (Figs. S21−S27), confirming its carbon skeleton as ergosterol-like.^29^ Up to now, all naturally occurring steroids from fungi have been found to exclusively possess a hydroxyl group at C-3. Surprisingly, the typical signals for hydroxylation at C-3 of ergosterol were absent in the 1D NMR data of the metabolite (**8**), which was eventually characterized as an oxygen-free ergosterene based on the NMR and HRMS data, (22E)-ergosta-3,5,7,9(11),22-pentaene (Figs. 5E and S28). The existence of ergosterene (**8**) in the mutant Δ*An* indicated hypoxia inside in Δ*An*, consistent with the decreased OCRs in *An*.

Up to now, no any oxygen free ergosterenes have been reported in fungi. Notably, oxygen-containing ergosterol is a well-known steroid crucial for maintaining cell membrane integrity and fluidity in fungal cell membrane.^29^ Meanwhile, ergosterol is also a component in secretory vesicles. We evaluated the contents of ergosterol and ergosterene between the mutant Δ*An* and WT at 37 °C. Quantitative analysis showed an astonishing accumulation of ergosterene **8** in Δ*An*, with 1943.5 times higher concentration than in WT (Figs. 5F−5G), while ergosterol contents were markedly decreased in Δ*An* compared to WT, suggesting that oxygen free ergosterene (**8**) should be the major steroid component in the fungal membrane and vesicles in Δ*An* at 37 °C. The distinct structure features of oxygen-free ergosterene (**8**) and ergosterol resulted in the distinct endocytosis and vesicles between Δ*An* and WT.

Interestingly, though addition of carviolin A increased the levels of ergosterol in both WT and the Δ*An* mutant compared with solvent, the level of oxygen-free ergosterene in the Δ*An* mutant were also slightly increased (Figs. 5F−5G). This result was consistent with the enlarged vesicles for sequestering carviolin A-induced granules in the Δ*An* mutant (Figs. 4g−4i). Additionally, carviolin A treatment greatly increased the levels of TTP derived oxygen-containing iron chelators in both WT and mutant Δ*An* (Figs. 5H−5K). Notably, the Δ*An* mutant treated with carviolin A could not produce as many ergosterol and iron chelators as WT, consistent with less granules in the Δ*An* mutant than those in WT. Moreover, the total levels of oxygen-free TTPs E−F (**5**−**6**) were much higher than those of oxygen-containing TTPs A−D (**1**−**4**) in the Δ*An* mutant, while the case was just opposite in WT (Figs. 5J−5K). All the results confirmed that the hypoxia inside in Δ*An* inhibited formation of the oxygen-containing metabolites.

### Evaluation of functions of biological active polycyclic aromatics in iron levels and ratios of fungal and human cells

We observed that carviolin A treatment significantly increased levels of Fe^2+^ and total free iron in the mycelia of both WT and Δ*An* (Figs. 6A−6B). Moreover, in the broths of Δ*An*, the levels of Fe^2+^, Fe^3+^ and total free iron were all decreased (Figs. 6C−6D). All the results suggested that exogenous carviolin A could induce Fe^2+^ influx and increase ratio of Fe^2+^/Fe^3+^ in fungal mycelia, which allowed us to propose that the cardiotoxicity of anthraquinones in treating cancers might be asribed to iron overload in cardiac cells.

**Figure 6.**
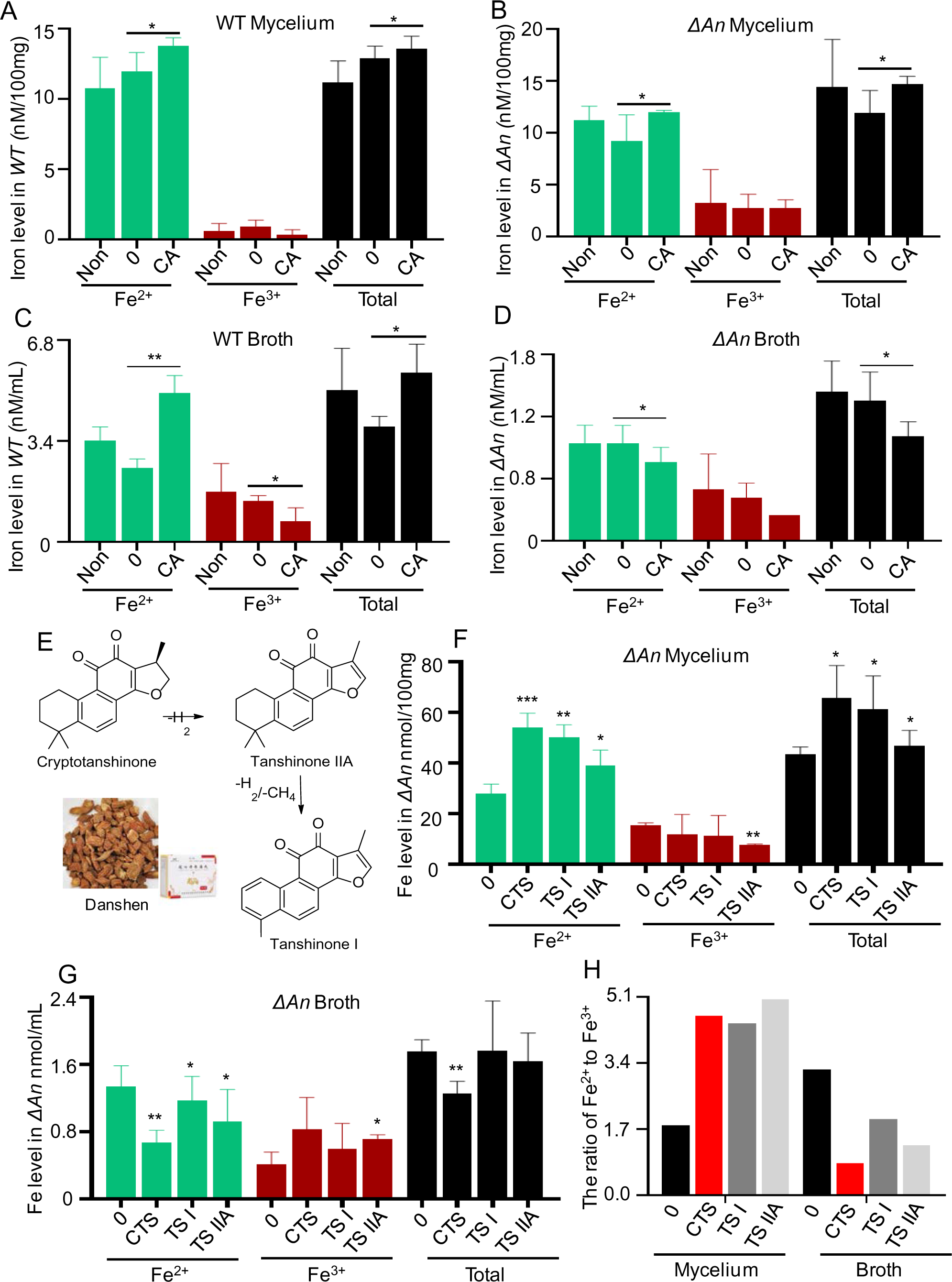
Evaluation of the roles of carviolin A (CA), tanshinone IIA (TS IIA), tanshinone I (TS I) and cryptotanshinone (CTS) in fungal iron transports and iron compositions. **A−D)** Comparison of iron levels in mycelia (A−B) and broths (C−D) of WT and the mutant Δ*An* treated with CA. Non: no treatment. 0: treatment with only solvent. E) Structures of three major components, TS IIA, TS I and CTS from Danshen and their biosynthetic pathways. F−G) The effects of TS IIA, TS I and CTS on iron levels in the mycelia (F) and broths (G) of the mutant Δ*An* at 37 °C. 0: treatment with only solvent. H) The effects of TS IIA, TS I and CTS on the ratios of Fe^2+^/Fe^3+^ in the mycelia and broths of the mutant Δ*An* at 37 °C. 0: treatment with solvent. p < 0.05 (*), p < 0.01 (**), p < 0.01 (***).

Danshen, the dried root of *Salvia miltiorhiza* Bunge (Lamiaceae), has been used for centuries in traditional Chinese medicine (TCM) for treating various symptoms such as cardiovascular and cerebrovascular diseases. In recent years, it has gained acceptance as a natural health product in Western countries.^30,31^encouraged us to investigate the iron transport-related functions of its main ingredients. A class of terpenoid-derived polycyclic aromatic metabolites, including tanshinone IIA (TS IIA), tanshinone I (TS I) and cryptotanshinone (CTS), are key medicinal active ingredients found in Danshen (or Tanshen), (Fig. 6E). We treated Δ*An* with 20 μM of these three metabolites at 37 °C. The negative control consisted of strain treated with only solvent DMSO. Remarkably, all the three plant metabolites significantly increased the levels of Fe^2+^ and total iron while decreasing Fe^3+^ levels in the mycelia of Δ*An*, compared with only solvent (Fig. 6F). Opposite results were observed in the broth of the mutant Δ*An* (Fig. 6G). The three plant metabolites all decreased the levels of Fe^2+^ and total iron compared with only solvent (Fig. 6G). All the three plant metabolites substantially increased the ratio of Fe^2+^/Fe^3+^ in the mycelia but decreased the ratio in the broths (Fig. 6H). These findings closely paralleled the effects observed with carviolin A treatment, suggesting that these three plant polycyclic aromatic metabolites, TS IIA, TS I and CTS, could induce membrane ferrous iron transport from the mycelia to the broths in the fungus.

To investigate anthraquinone cardiotoxicity, we evaluated the effects of anticancer agent mitoxantrone (Mito) on the levels of Fe^2+^ and total iron in human hepatocellular carcinomas cells (HepG2) and carviolin A, TS IIA, TS I and CTS were used as positive controls. Notably, Mito and TS IIA displayed strong inhibitory activity against HepG2 cells compared with solvent (Fig. 7A) and others exhibited no significant inhibition. Moreover, Mito demonstrated stronger anticancer activity than TS IIA. Interestingly, Mito and TS IIA, together with carviolin A, caused strongly increased levels of Fe^2+^ and total iron in HepG2 cells, compared to solvent (Fig. 7B). In particular, Mito treatment caused the highest levels of Fe^2+^ and total iron in HepG2 cells, and TS IIA seconded. The ratios of Fe^2+^/Fe^3+^ in HepG2 cells treated with Mito, TS IIA, and carviolin A, were also sharply elevated compared with solvent (Fig. 7C).

**Figure 7.**
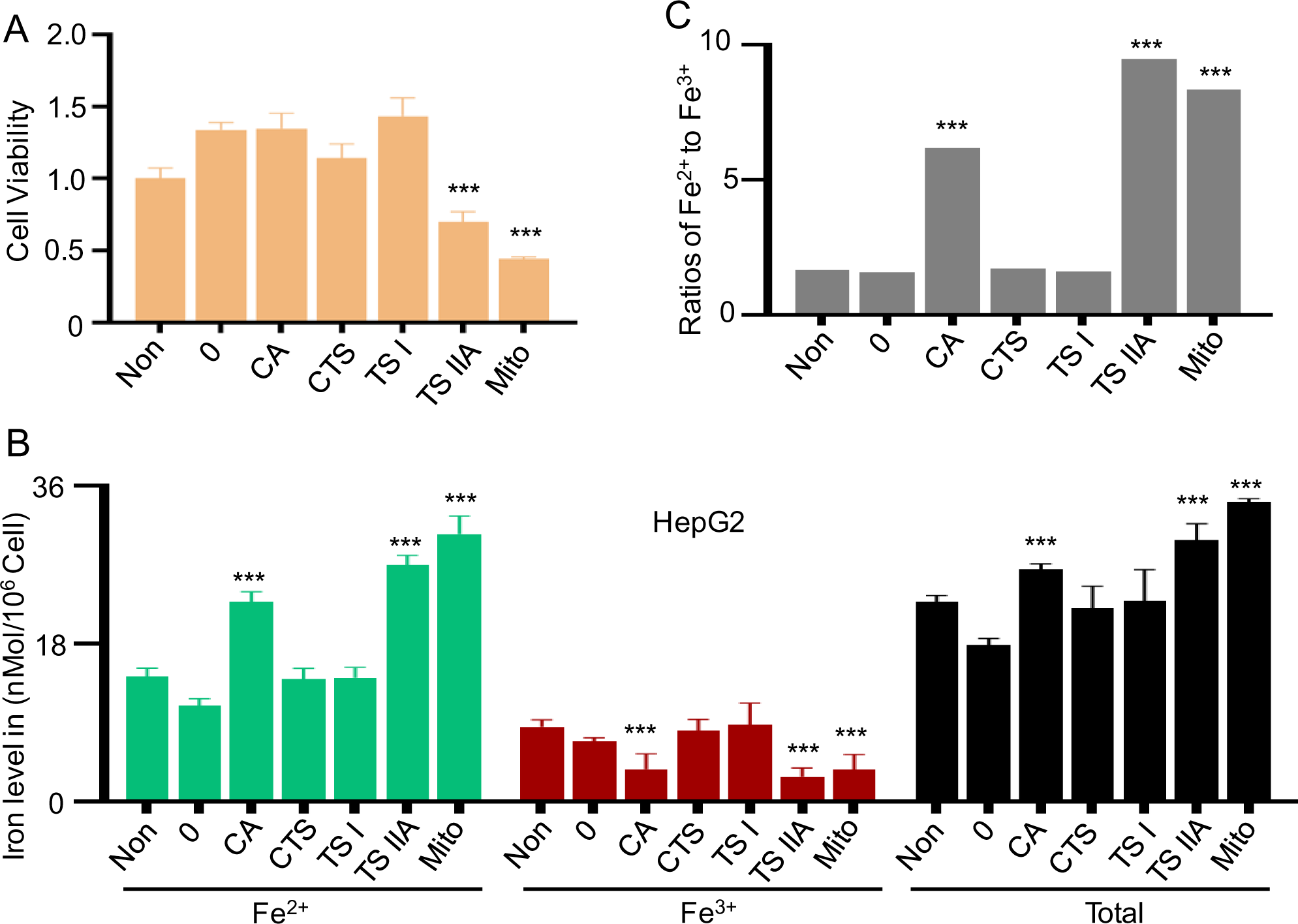
Evaluation of the roles of carviolin A (CA), tanshinone IIA (TS IIA), tanshinone I (TS I), cryptotanshinone (CTS) and mitoxantrone (Mito) in cell viability (A), and iron levels (B) and ratios (C) of HepG2 cells.

## DISCUSSION

Temperature variations induce metabolic changes in organisms to meet specific energy requirements.^32−35^ Iron metabolism is crucial for energy production, metabolite synthesis, and oxygen transportation.^36–41^ Deciphering how organisms regulate iron homeostasis and metabolism in response to temperature changes is pivotal in understanding their community specificity and growth preference.^42^ However, information about the effect of temperature on iron homeostasis and metabolism remains limited. In this study, we found that *T. dupontii* WT accumulated PKS-derived anthraquinone pigments in response to cold stress. In particular, *T. dupontii* mutant Δ*TTP* with lack of PIAs derived iron chelators dramatically produced much more anthraquinones than WT. Disruption of the *PKS* gene *An* for the anthraquinones biosynthesis not only caused loss of anthraquinones but also sharply decreased levels of PIA derived iron chelators in mutant Δ*An*. Interestingly, Δ*TTP* was enriched in anthraquinones while Δ*An* was deficient in anthraquinones. Previous study reported that Δ*TTP* displayed increased conidial formation,^11^ which was in sharp contrast to sharply decreased conidial formation in Δ*An* in the study. Importantly, mutant Δ*An* harbored more Fe^2+^ and total free iron in mycelia than WT while less in broths, which was in distinct contrast to Δ*TTP* that exhibited less Fe^2+^ and total free iron in mycelia than WT but more in broths. These results suggest that anthraquinones are involved in exporting Fe^2+^ from mycelia to broths and enhancing Fe^2+^/ Fe^3+^ ratios in mycelia.

Previous studies reported that redox cycles involving quinones occur through one-electron transfer, reducing Fe^3+^ to Fe^2+^, via secreted fungal hydroquinones and/or catechols,^27,43^ and some brown rot wood-decaying fungi and the ectomycorrhizal fungus *Paxillus involutus* have been shown to utilize Fenton chemistry via hydroquinones like metabolites to decompose organic matter.^43−45^ However, the underlying mechanisms on how these fungi use metabolites to manipulate Fenton chemistry have not been reported. Ferrous and oxygen are fundamental constituents for Fenton reactions, serving as a major source of oxidative power, a vital process in biology.^46^ Our findings indicate that anthraquinones not only reduce Fe^3+^ to Fe^2+^, but also excrete Fe^2+^ from fungal mycelia to ambient environment, which make the Fenton reactions feasible to decompose organic matters and elevate ambient temperatures.

We observed that the mutant Δ*An* contained less intracellar black granules in mycelia and extrudes in broths than WT, besides having few iron chelators and no anthraquinones. However, Δ*An* displayed inner-membrane derived vesicles while WT not. Supplement bioassay indicated that exogenous carviolin A, a major anthraquinone component, could induce extracellar granule formation and import to both Δ*An* and WT. In contrast to WT into which granules easily entered, Δ*An* used the vesicles to sequester the exogeous granule from the inner in mycelia, suggesting that Δ*An* adopted an unprecedented lifestyle with shutdown membrane transport through transcriptional analysis, we found that steroid-mediated membrane transport was the most down-regulated. Eventually, metabolic analysis and detailed chemical investigation led to characterization of a rare nature-occurring oxygen free ergosterene the mutant Δ*An*, consistent with the dramatically decreased formation of oxygen-containing metabolites, such as TTP-derived iron chelators and ergosterols in Δ*An.* All the results suggested that loss of anthraquinones caused the mutant Δ*An* to adopt a strategy of isolation via sharply decreasing the endocytosis and applying the ergosterene -derived vesicles to sequester exogenous granules induced by carviolin A, thus leading to hypoxia inside.

Up to now, most studies suggested that sterol compositions influence endocytosis, such as defective in the sterol-biosynthesis mutant *cyclopropylsterol isomerase1-1* (*cpi1-1*) displayed altered membrane sterol composition, leading to endocytosis-defective *cpi1-1* cells.^47,48^ However, no oxygen-free sterenes have been found to be related to endocytosis inhibition. Not to mention that anthraquinones biosynthesis could improve endocytosis via enhancing oxygen-dependent sterol biosynthesis.^49,50^ Our finding suggested the functions of anthraquinones biosynthesis in controlling sterene derived sterol biosynthesis and thus membrane transports. Previous studies indicated that anthraquinone biosynthesis involved oxygen consumption and carbon dioxide emission,^36^ which are in favor of Fe^2+^, instead of Fe^3+^. Moreover, the anthraquinone reduction of Fe^3+^ to Fe^2+^ and thus the increase in the Fe^2+^/Fe^3+^ ratios might booster Fe^2+^ efflux with anthraquinone excretion from fungal mycelia.

Previous studies on the repertoire conferring iron homeostasis in organisms have highlighted the functional collaboration of multiple proteins in iron transports, iron metabolism, iron storage and related signal pathways.^36–42^ Up to date, no physiologic mechanisms for iron excretion have been characterized yet in fungi, plants and animals, only, the mechanisms for iron absorption and storage have been reported to maintain iron homeostasis.^42^ In this study, we found that a thermophilic fungus *T. dupontii* with a notably reduced set of genes,^11^ used polycyclic aromatic metabolites to dictate Fe^2+^ efflux, steroid-mediated endocytosis, and iron-containing granule formation. The conventional array of genes for absorption, storage and excretion to control iron metabolism and homeostasis might be inadequate for enabling the fungus to thrive in response to temperature reduction. The thermophilic fungus has to adopt a most concise and quick tactic to regulate iron metabolism in response to temperature fluctuations since the biosphere temperatures are commonly unsuitable for thermophilic fungal growth. Notably, only three genes are required for the carviolin A formation. This fungal adaption strategy proves to be effective and efficient, as evidenced by the fact that *T. dupontii* is a dominant species among thermophilic fungi.

The above results allowed us to propose that iron overload in cardiac cells induced by anthraquinones-mediated Fe^2+^ transport might cause the outstanding cardiotoxicity of anthraquinones in cancer treatment. Interestingly, the major components in medical plant Danshen for treating cardiovascular diseases for centuries in China, belong to terpenoid-derived polycyclic aromatic metabolites. Here, we found that these components also demonstrated potent ferrous transporting capabilities. These findings shed new light on pharmacological mechanisms of anthraquinones and other types of polycyclic aromatic metabolites in fungi, and medical plants and herbs. These polycyclic aromatic metabolites may be new potential candidates for iron-homeostasis therapeutics and drug delivery.

## Acknowledgment

We would like to thank the Kunming Biological Diversity Regional Center of Instrument, Kunming Institute of Zoology, Chinese Academy of Science for help with transmission electron microscopy. We are grateful to Guo Yingqi and Wu Xingcai for their great help in preparing TEM samples and taking TEM images, and to Joan Wennstrom Bennett for manuscript editing.

