## Supporting information for "Thermophilic Fungus Uses Anthraquinones to Modulate Ferrous Excretion, Sterol-Mediated Endocytosis and Iron Storage in Response to Cold Stress"

### Anthraquinone-Mediated Modulation of Ferrous Excretion, Sterol-Mediated Endocytosis and Iron Chelators in Thermophilic Fungal Response to Cold Stress

Shuhong Li,[1,⊥] Donglou Wang,[1,⊥] Jiangbo He,[2,⊥] Chunhua Liao,[1] Zhangxin Zuo,[1] Shenghong Li,[3] Xuemei Niu[1,*]

1 State Key Laboratory for Conservation and Utilization of Bio-Resources & Key Laboratory for Microbial Resources of the Ministry of Education, School of life Sciences, Yunnan University, Kunming 650091, P. R. China

2 Kunming Key Laboratory of Respiratory Disease, Kunming University, Kunming 650214, P. R. China

3 State Key Laboratory of Phytochemistry and Plant Resources in West China, Kunming Institute of Botany, Chinese Academy of Sciences, Kunming 650201, P. R. China

Running head: Anthraquinone-Mediated ferrous excretion, iron storage and endocytosis

[⊥]These authors contributed equally.

**Supplementary Tables**
**Table S1.** Primers used for construction of the mutant Δ*An* (Δ*PKS30*).

**Supplementary Figures**
**Fig. S1.** A class of tryptophanderived cyclodipeptides with one or two prenyl groups, talathermophilins A–F (TTPs A–F, **1–6**), were remarkably elevated in WT at 37 °C vs. 45 °C.
**Fig. S2.** The HRMS of anthraquinone metabolite carviolin A (**7**).
**Fig. S3.** The UV absorptions of anthraquinone metabolite carviolin A (**7**).
**Fig. S4.** The 1HNMR spectrum of anthraquinone metabolite carviolin A (**7**).
**Fig. S5.** The $^{13}$CNMR spectrum of anthraquinone metabolite carviolin A (7).
**Fig. S6.** Quantitative analysis focuses on metabolite **7**, known as carviolin A, in the fungal Δ*TTP* vs. WT at 37 °C
**Fig. S7.** A series of anthraquinone metabolites, including 4 anthraquinone dimmers and 5 monomers, were remarkably elevated in Δ*TTP* vs. WT at 37 °C.
**Fig. S8.** The effect of carviolin A on ferric reduction.
**Fig. S9.** The *An* gene was among the most significantly up-regulated core biosynthetic genes in the fungal WT at 37 °C vs. 45 °C.
**Fig. S10.** Schematic of homologous recombination of Δ*An* (Δ*PKS30*); Confirmation of the Δ*An* mutants by PCR.
**Fig. S11.** Quantitative analysis of carviolin A contents in Δ*An* vs. WT at 37 °C and 45 °C.
**Fig. S12.** A–B) Quantitative analysis of superoxide ROS levels in Δ*An* and WT at 37 °C and 45 °C. C–D) Analysis of the conidiation (C) and spore germination rate (D) in WT and the Δ*An* mutant at 37 °C and 45 °C.
**Fig. S13.** Quantitative analysis of the TTPs A–F in Δ*An* and WT for 7days.
**Fig. S14.** Quantitative analysis of the key genes in *An* gene cluster and *TTP* gene in Δ*An* and WT.
**Fig. S15.** The effect of anthraquinone on $Fe^{2+}$ efflux.
**Fig. S16.** The colony growths of Δ*An* and WT on Congo red.
**Fig. S17.** Quantitative analysis of black granules in each cell at 37 °C and 45 °C.
**Fig. S18.** Quantitative analysis of black granules in each cell by chemical complementation with carviolin A (7).
**Fig. S19.** Comparative analysis of endocytosis between WT and the Δ*An* mutant using the chemical probe for $Fe^{2+}$.
**Fig. S20**. KEGG analysis of transcriptional profiles revealed the down-regulated (A) and up-regulated (B) pathways in Δ*An* vs. WT.
**Fig. S21.** Mass spectrum of ergosterene (**8**).
**Fig. S22.** $^{1}$H NMR spectrum of ergosterene (**8**).
**Fig. S23.** $^{13}$C NMR spectrum of ergosterene (**8**).
**Fig. S24.** HSQC spectrum of ergosterene (**8**).
**Fig. S25.** $^{1}$H-$^{1}$H COSY spectrum of ergosterene (**8**).
**Fig. S26.** HMBC spectrum of ergosterene (**8**).
**Fig. S27.** ROESY spectrum of ergosterene (**8**).
**Fig. S28.** Structures of ergosterene (**8**) and ergosterol.

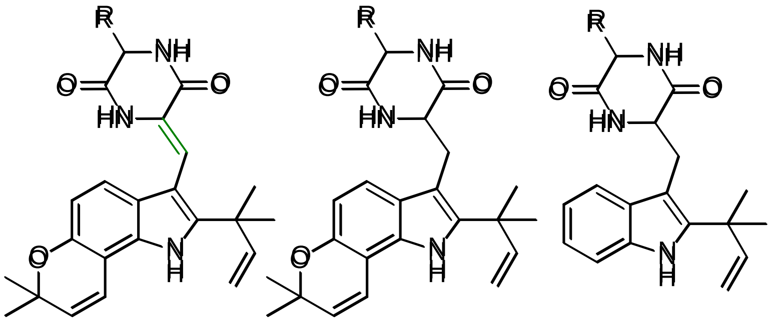

(**1**) R = H
(**2**) R = Me

(**3**) R = H
(**4**) R = Me

(**5**) R = H
(**6**) R = Me

**Figure S1.** A class of tryptophanderived cyclodipeptides with one or two prenyl groups, talathermophilins A–F (TTPs A–F, **1**–**6**), were remarkably elevated in WT at 37 °C vs. 45 °C.

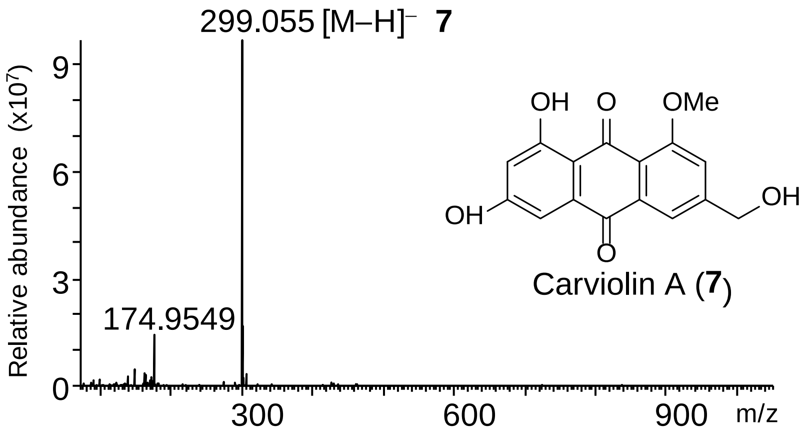


**Figure S2.** The HRMS of anthraquinone metabolite carviolin A (**7**).

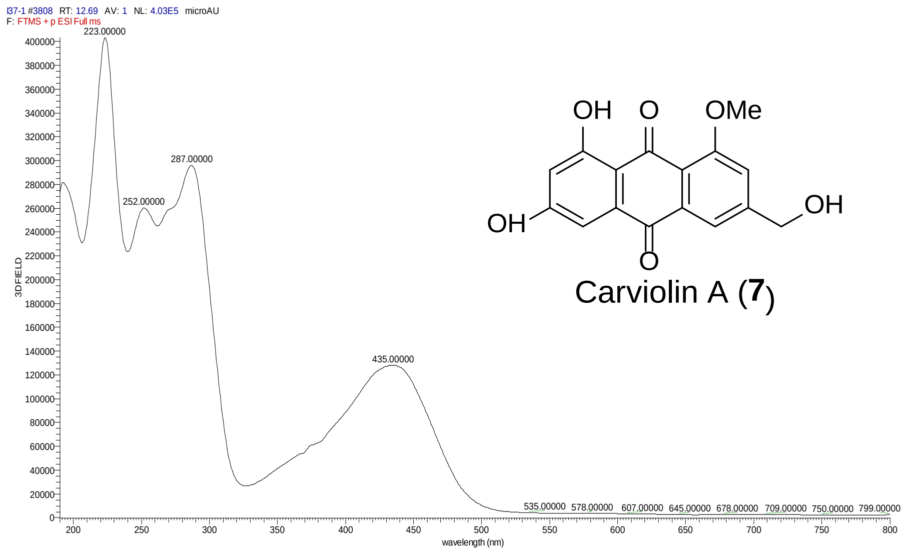


**Figure S3.** The UV absorptions of anthraquinone metabolite carviolin A (**7**).

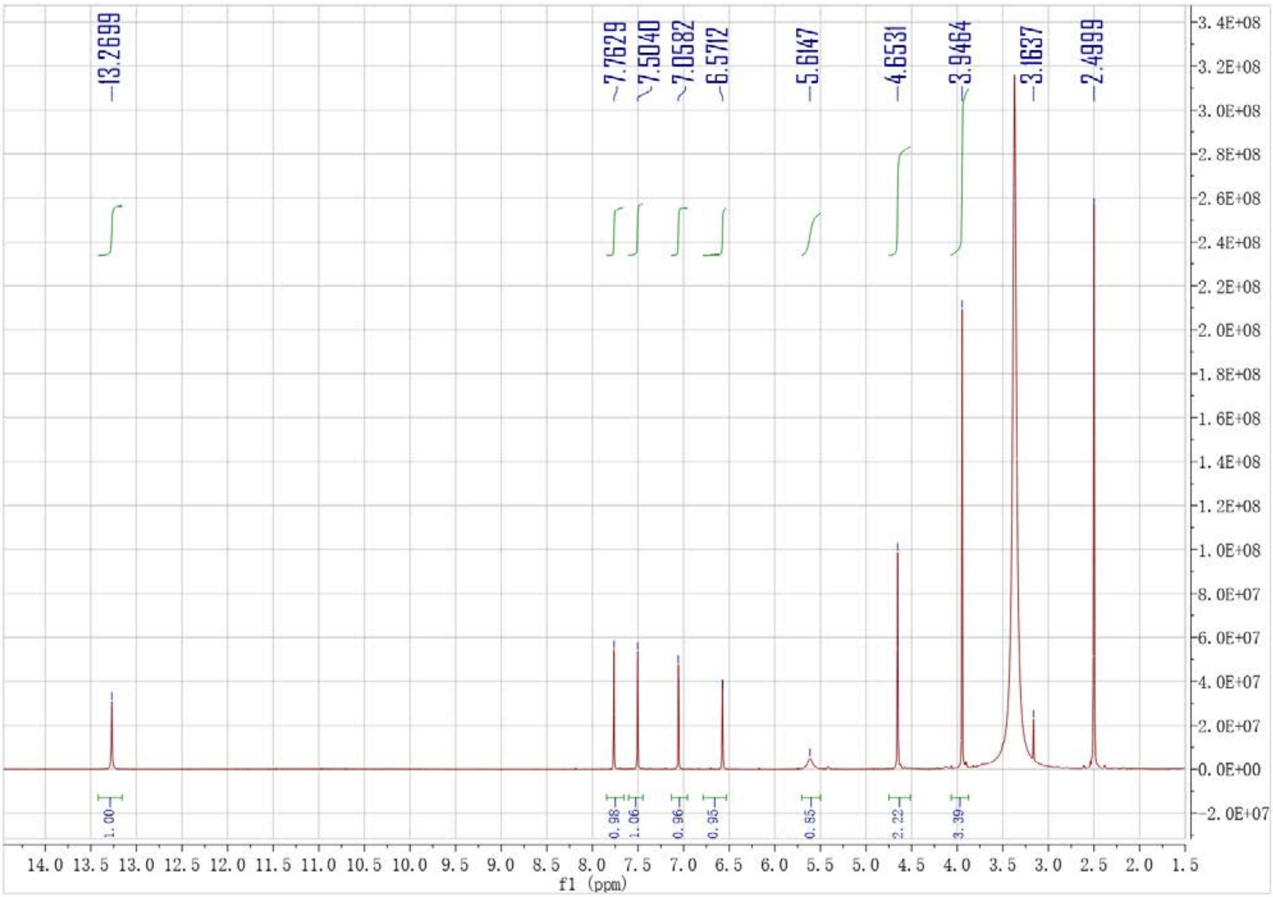


**Figure S4.** The $^{1}$HNMR spectrum of anthraquinone metabolite carviolin A (**7**).

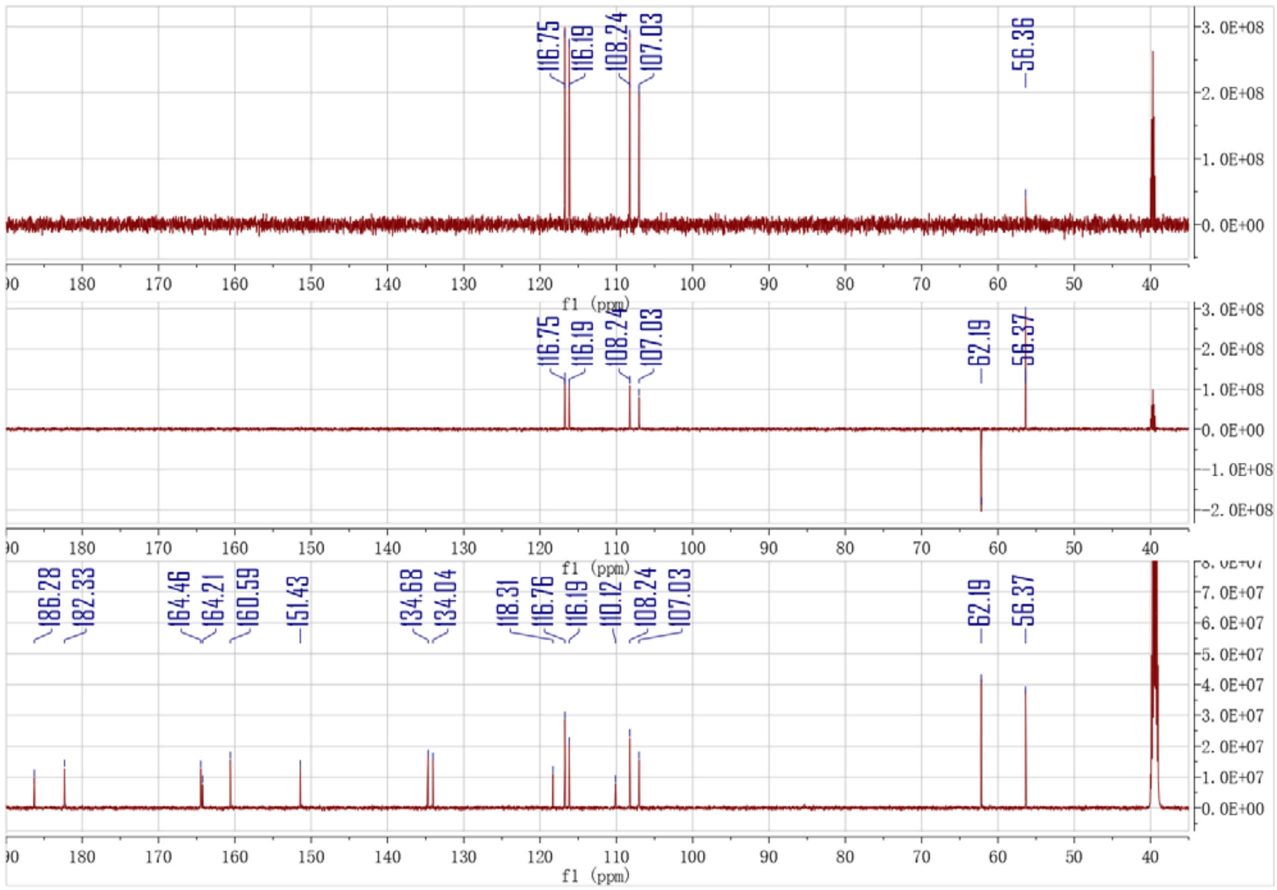


**Figure S5.** The $^{13}$CNMR spectrum of anthraquinone metabolite carviolin A (**7**).

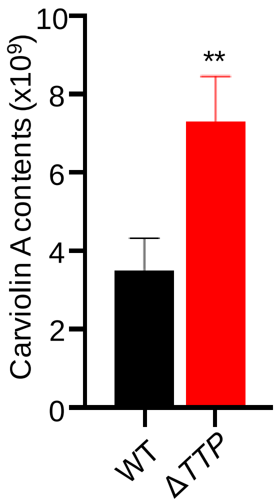


Figure S6 Quantitative analysis of carviolin A contents in Δ*TTP* vs. WT at 37 °C.

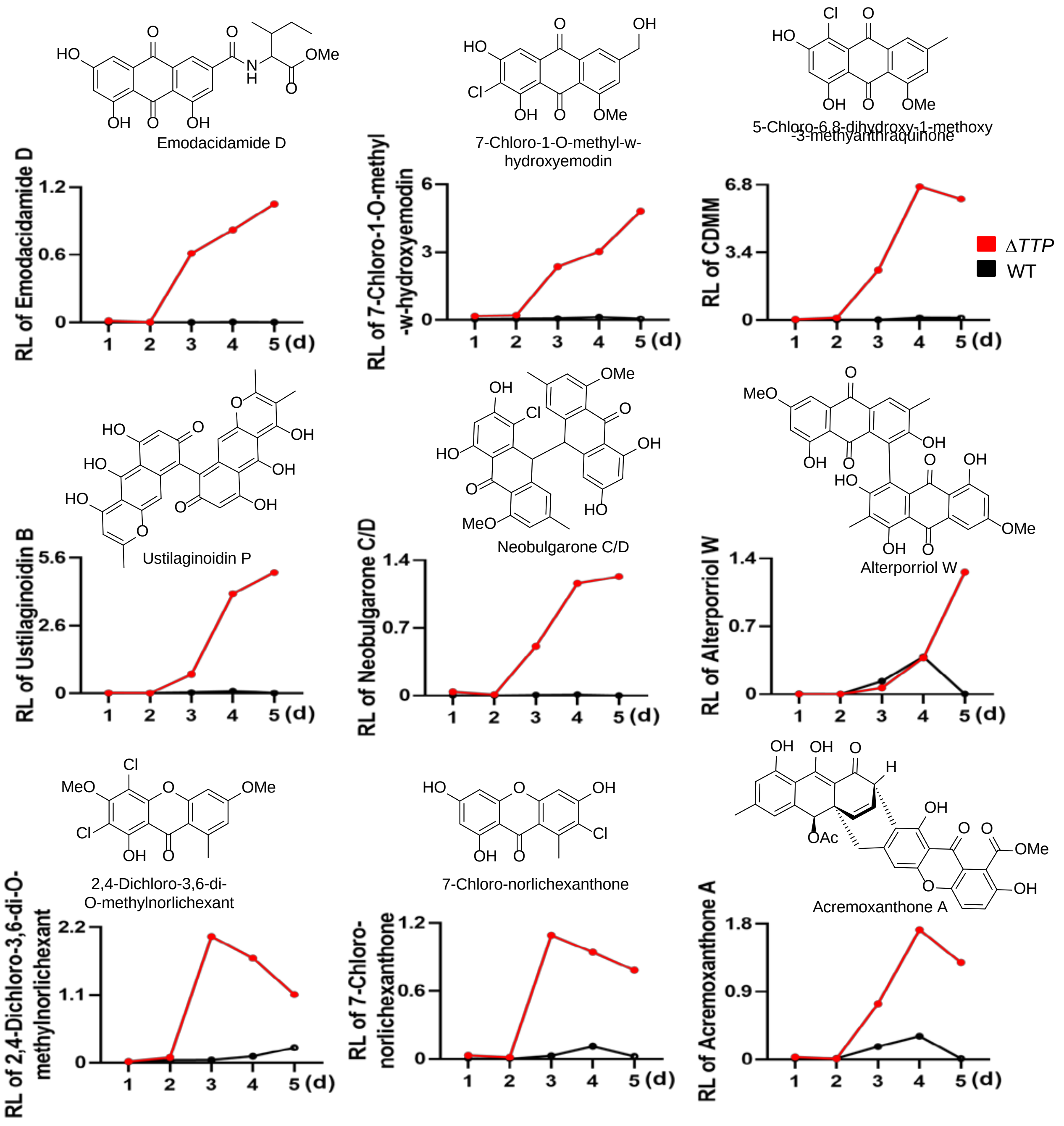


**Figure S7.** A series of anthraquinone metabolites, including 4 anthraquinone dimmers and 5 monomers, were remarkably elevated in Δ*TTP* vs. WT at 37 °C.

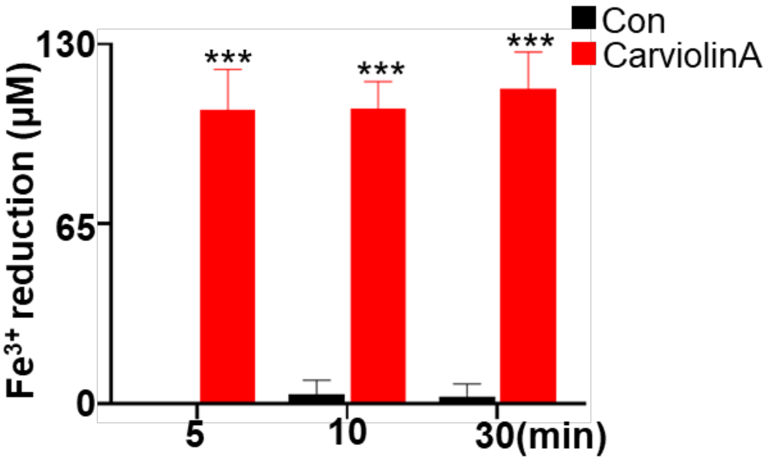


**Figure S8.** The effect of carviolin A on ferric reduction.

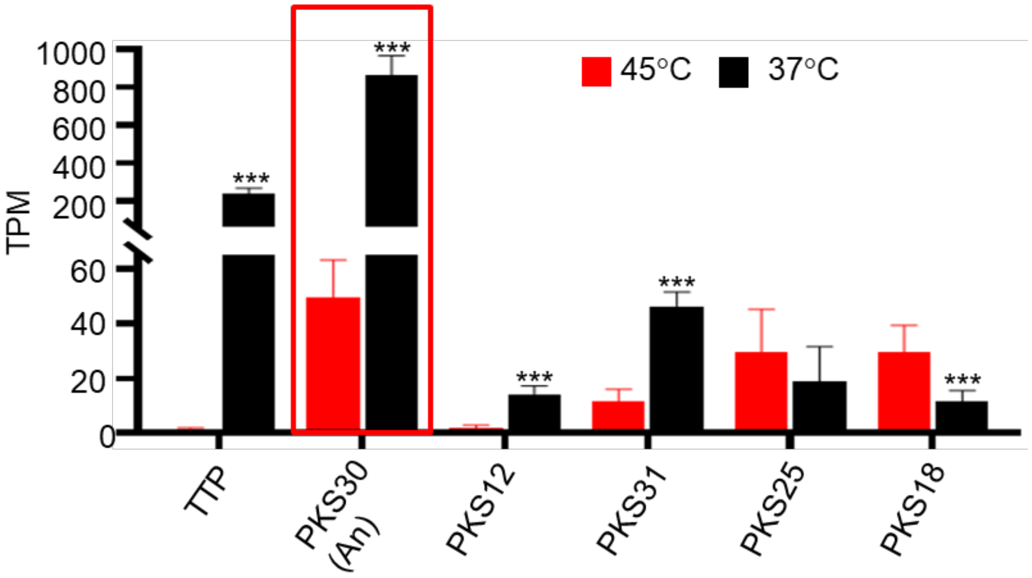


**Figure S9.** The *An* gene was among the most significantly up-regulated core biosynthetic genes in the fungal WT at 37 °C vs. 45 °C.

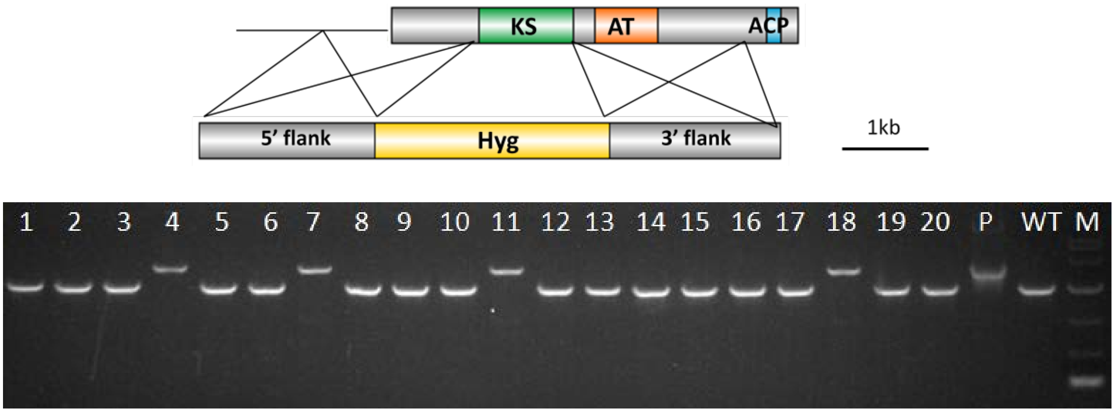


**Figure S10.** Schematic of homologous recombination of Δ*An* (Δ*PKS30*); Confirmation of the Δ*An* mutants by PCR. Line 4, 7, 11, 18: genomic DNA from the Δ*An* mutants. P: plasmid DNA as positive control; WT: genomic DNA from wild-type as negative control; M: DL 5000 DNA Marker.

**Table S1** Primers used for construction of the mutant Δ*An* (Δ*PKS30*).

| Primer name | Sequence (5' to 3') |
|---|---|
| PKS30 up F | AGTGCCAAGCTTGCATGCCTGCAGGTGCAGGCTGATGCGATCTATGGATC |
| PKS30 up R | CTTCTGTCGAGGCCTGATCATCGATGATGCCAGCAGTCAGGTAGTCGAG |
| PKS30-Hyg F | ATCTCGACTACCTGACTGCTGGCATCATCGATGATCAGGCCTCGACAG |
| PKS30-Hyg R | TTGAGCTCGCTCTTGATGCCGACATTCGGGGGATCCTCTAGATCTCGAC |
| PKS30 dw F | CGTCGAGATCTAGAGGATCCCCCGAATGTCGGCATCAAGAGCGAGCTC |
| PKS30 dw R | TCTGGTAAGCTTTGACCTCCTCGAGTCCACTTGTACGTCGGCAGAGTC |
| PKS30 YZ F | ACCAAGGTGTTCATCAGCGC |
| PKS30 YZ R | AGCCTCGATGTTGCCCTTC |

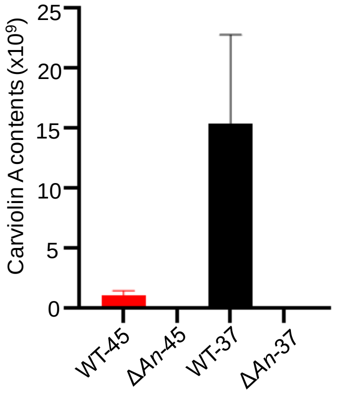


Figure S11 Quantitative analysis of carviolin A contents in Δ*An* vs. WT at 37 °C and 45 °C.

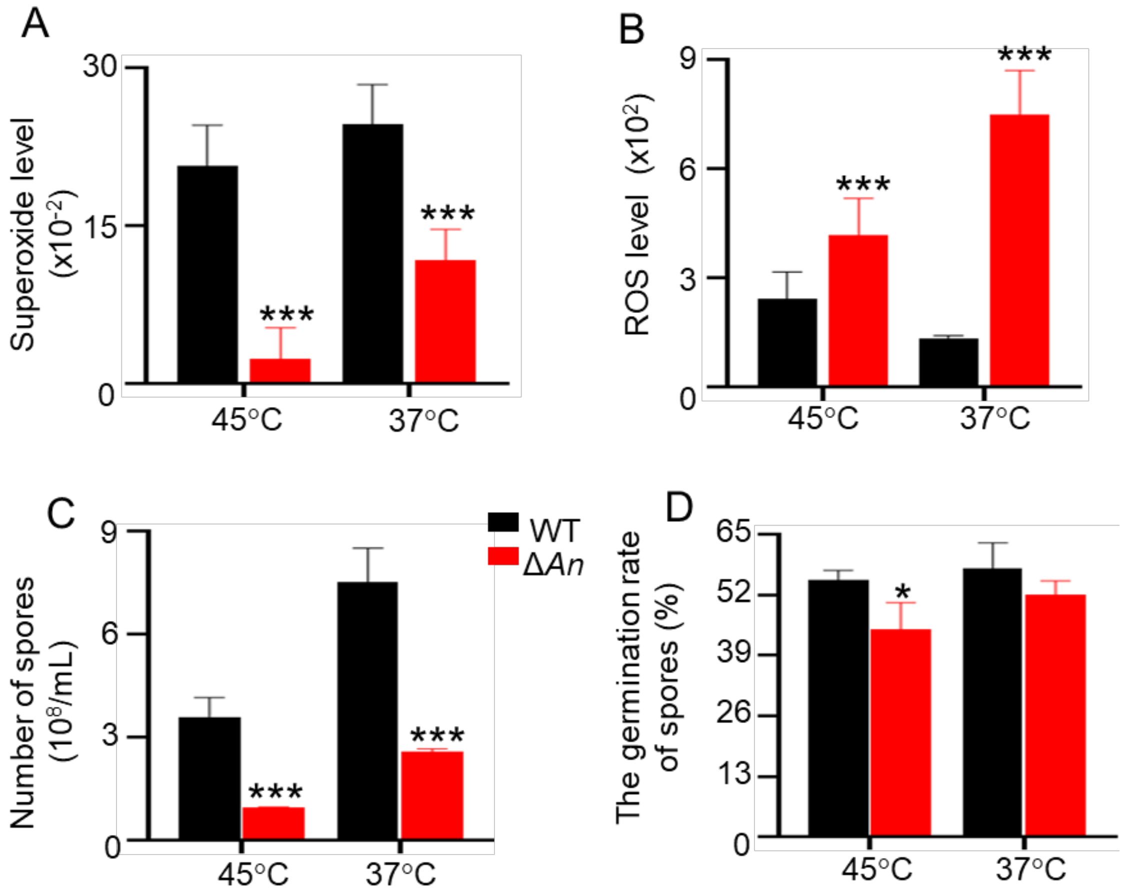


**Figure S12.** A–B) Quantitative analysis of superoxide ROS levels in Δ*An* and WT at 37 °C and 45 °C. C–D) Analysis of the conidiation (C) and spore germination rate (D) in WT and the Δ*An* mutant at 37 °C and 45 °C.

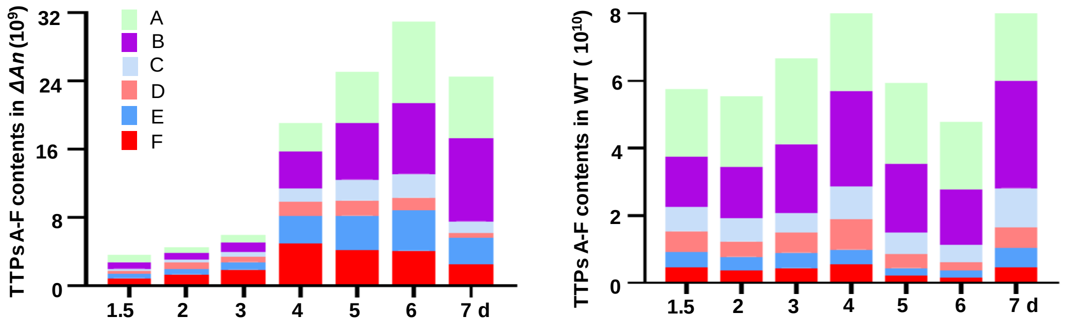


**Figure S13.** Quantitative analysis of the TTPs A–F in Δ*An* and WT for 7days.

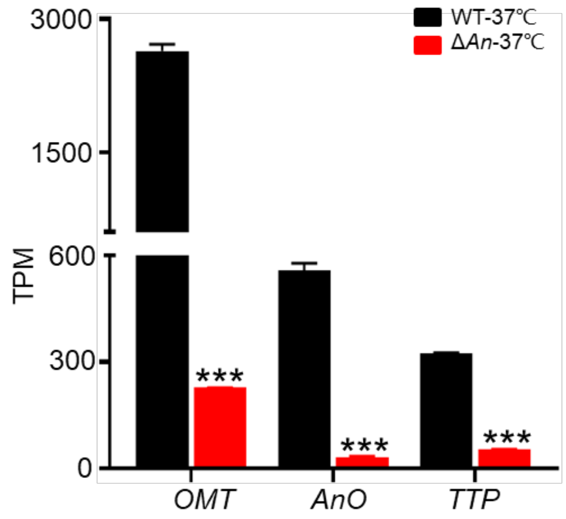


**Figure S14.** Quantitative analysis of the key genes in *An* gene cluster and *TTP* gene in Δ*An* and WT.

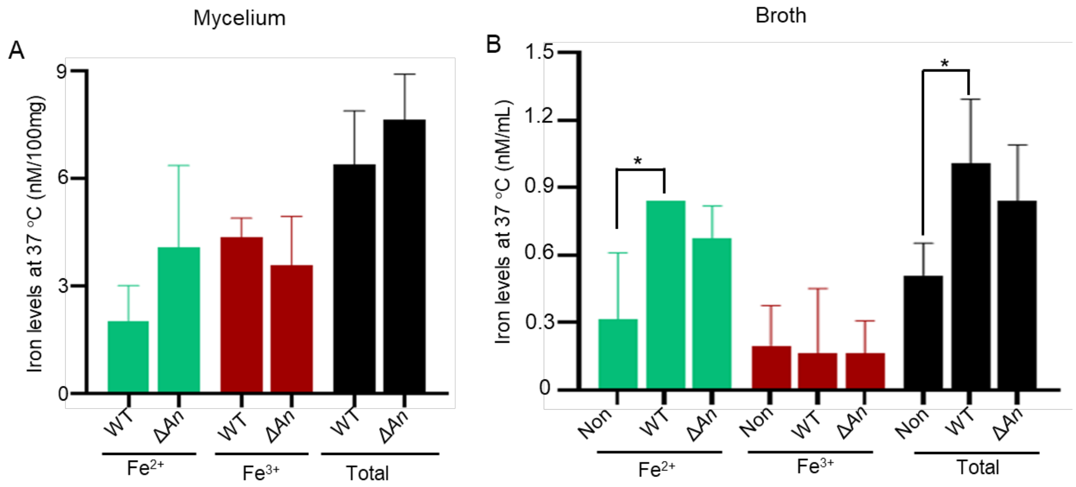


Figure S15. The effect of anthraquinones on $Fe^{2+}$ efflux. A: Comparison of the iron levels in mycelia of Δ*An* and WT grown on PDA with 0.1 mM $FeSO_4$ for 4 days at 37 °C; B: Comparison of the iron levels in broths of Δ*An* and WT for 2 days at 37 °C. Non: medium without fungal strains. *, *P* < 0.05; ***, *P* < 0.001.

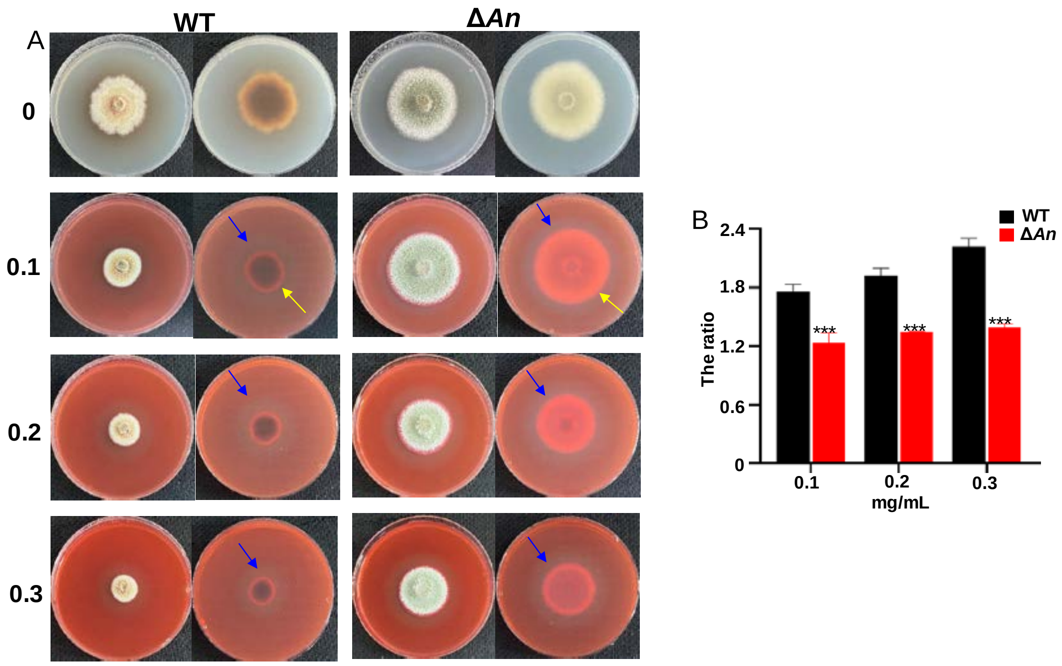


Figure S16. The colony growths of Δ*An* and WT on PDA with Congo red at different concentrations of 0.1-0.3 mg/mL at 37 °C. blue arrows: acidic zones of colonies; yellow arrows: fungal colonies; The ratios refer to the areas of acidic zones to the areas of fungal colonies. ***: $P < 0.001$.

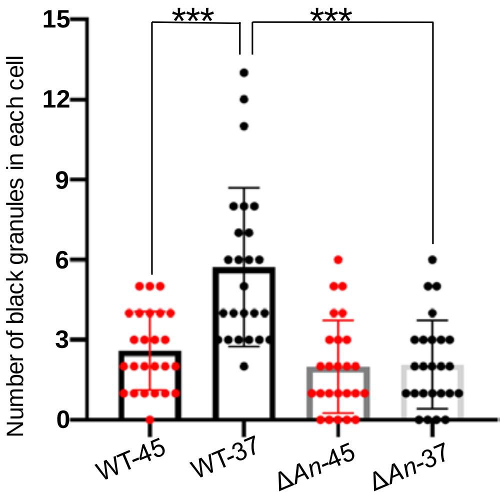


Figure S17. Quantitative analysis of black granules in each cell at 37 °C and 45 °C.

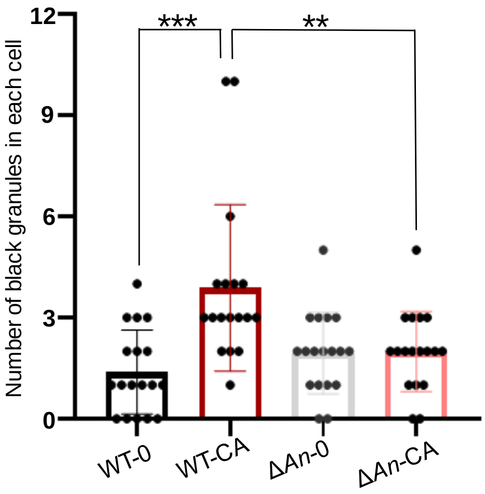


Figure S18. Quantitative analysis of black granules in each cell by chemical complementation with carviolin A (7).

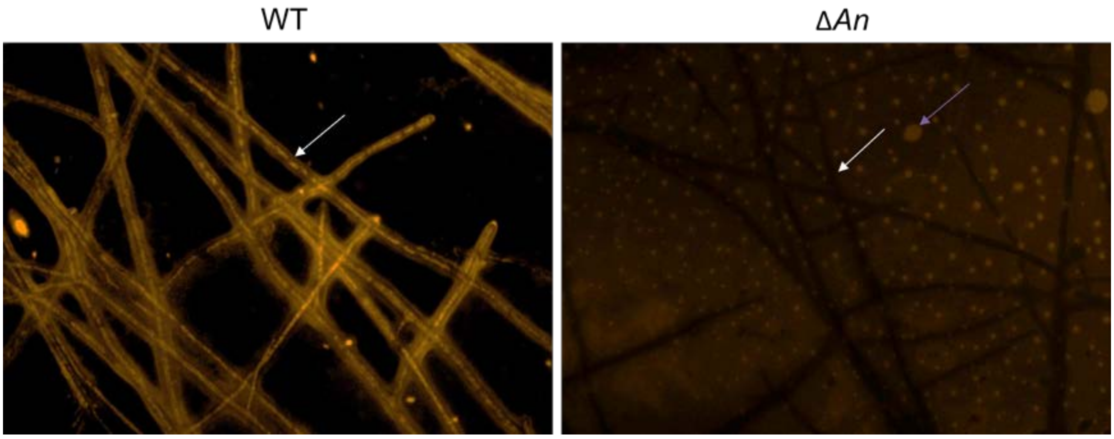


Figure S19. Comparative analysis of endocytosis between WT and the Δ*An* mutant using the chemical probe for $Fe^{2+}$.

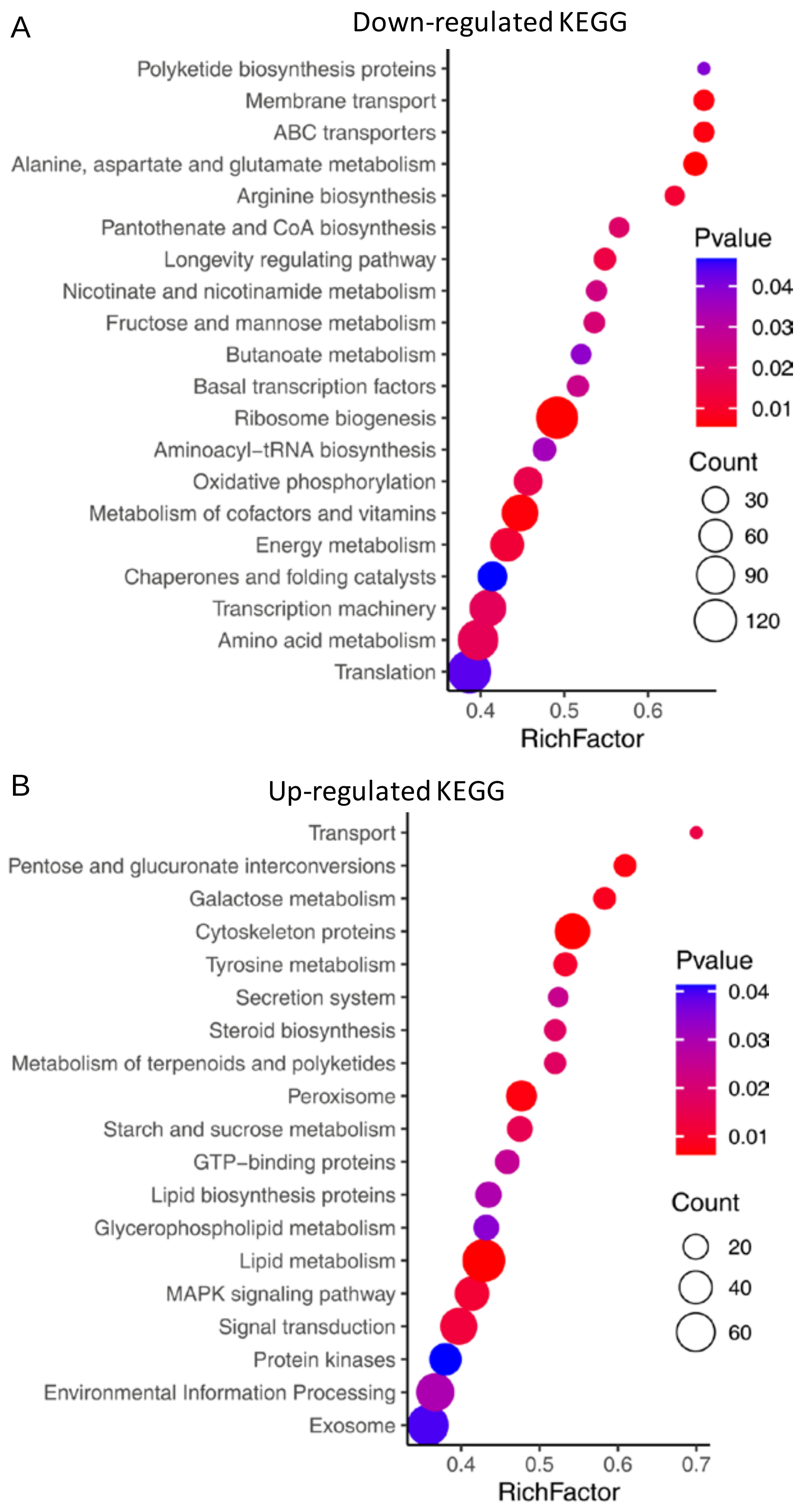


**Figure S20**. KEGG analysis of transcriptional profiles revealed the down-regulated (A) and up-regulated (B) pathways in Δ*An* vs. WT.

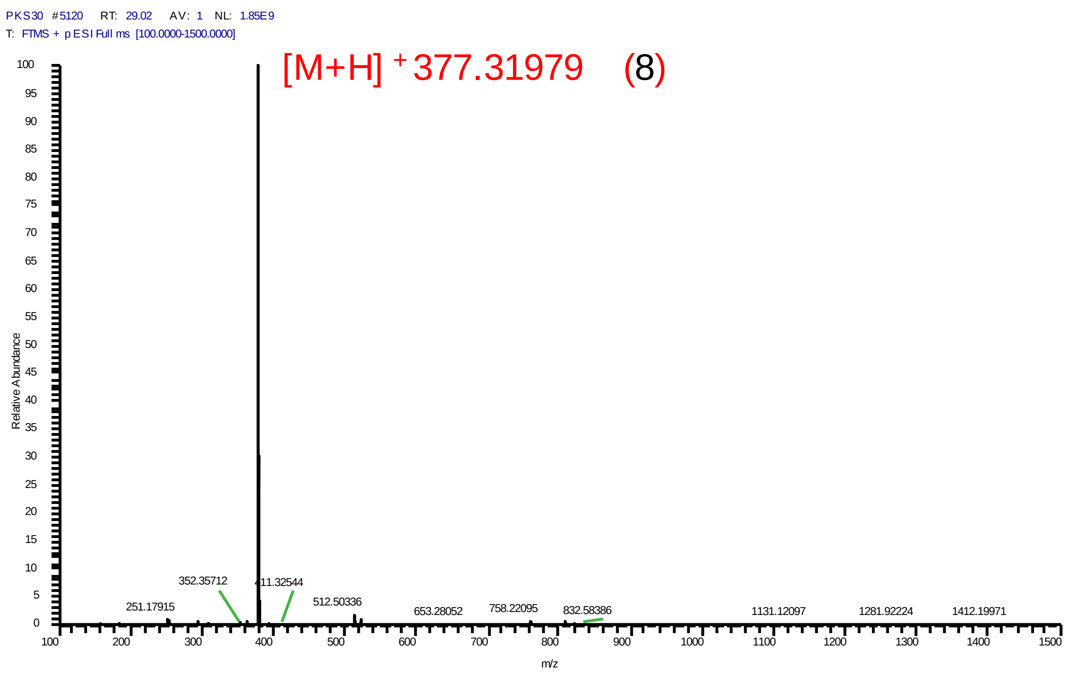


**Figure S21**. Mass spectrum of ergosterene (**8**).

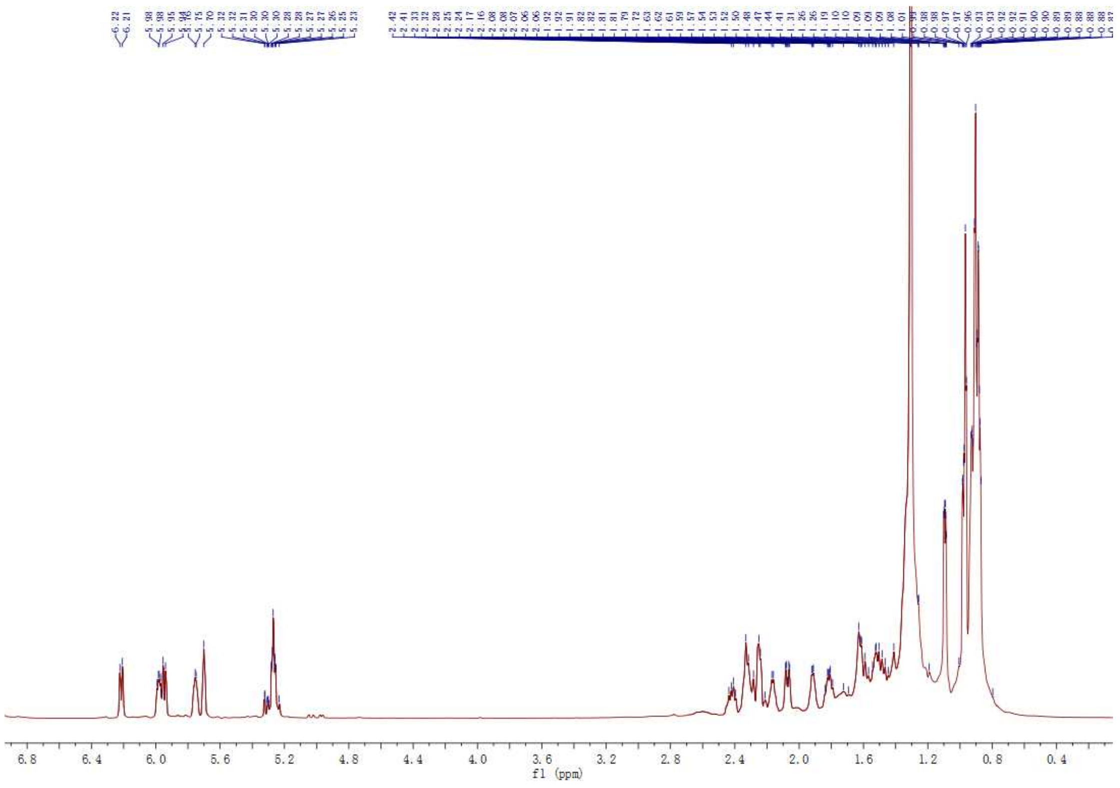


**Figure S22**. $^{1}H$ NMR spectrum of ergosterene (**8**).

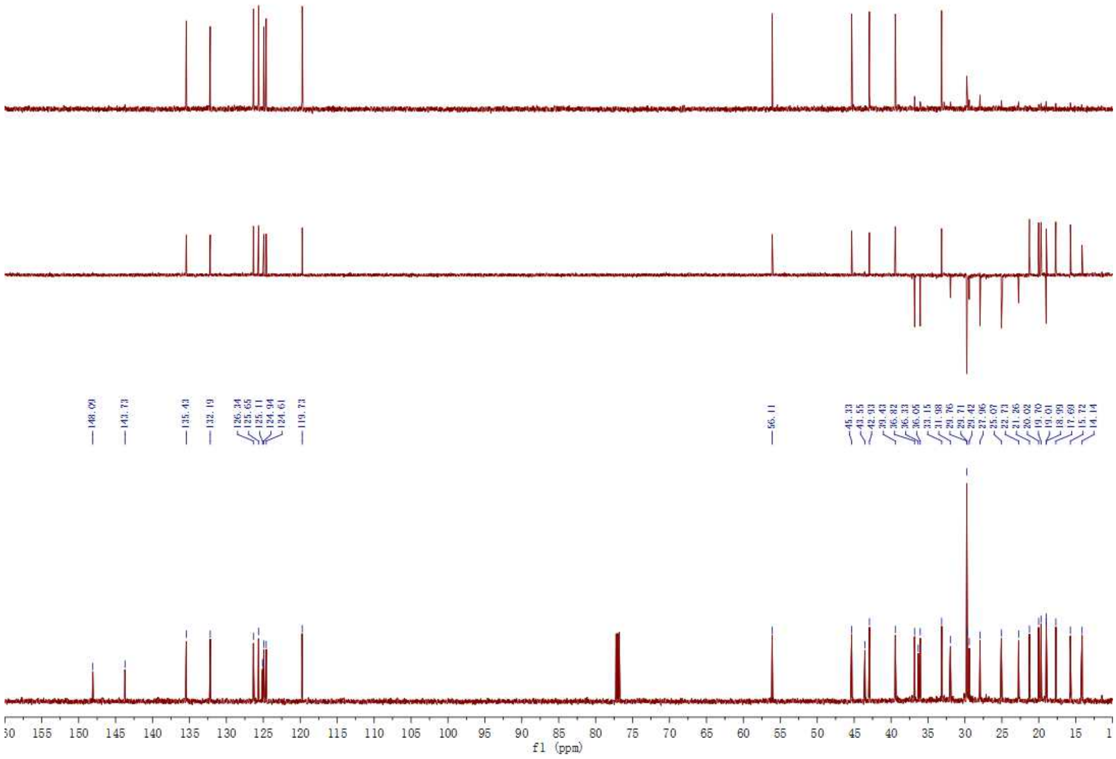


**Figure S23**. $^{13}C$ NMR spectrum of ergosterene (**8**).

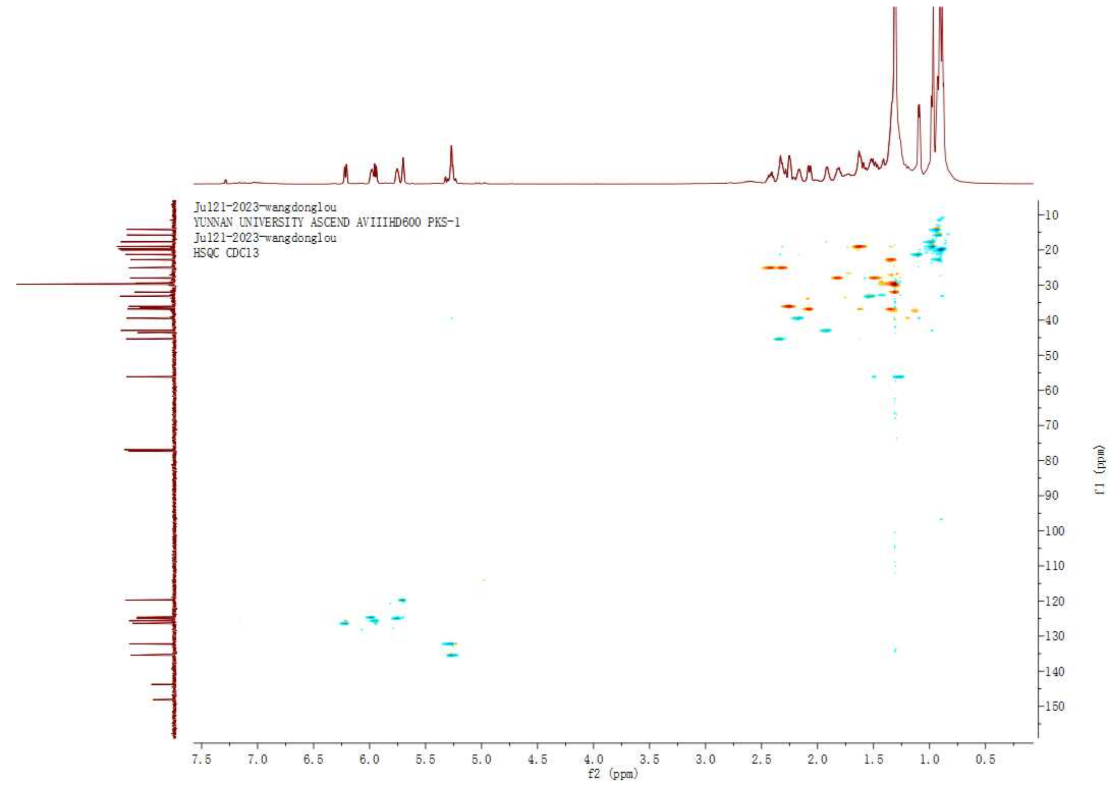


**Figure S24**. HSQC spectrum of ergosterene (**8**).

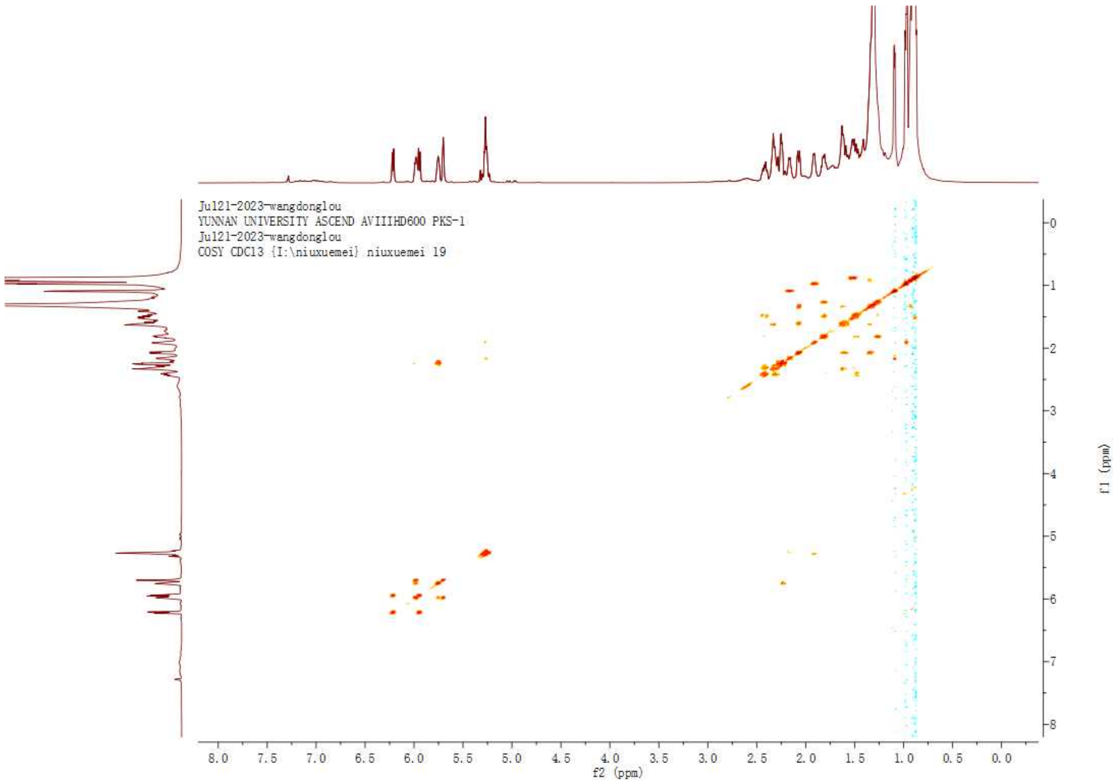


**Figure S25**. $^{1}H$-$^{1}H$ COSY spectrum of ergosterene (**8**).

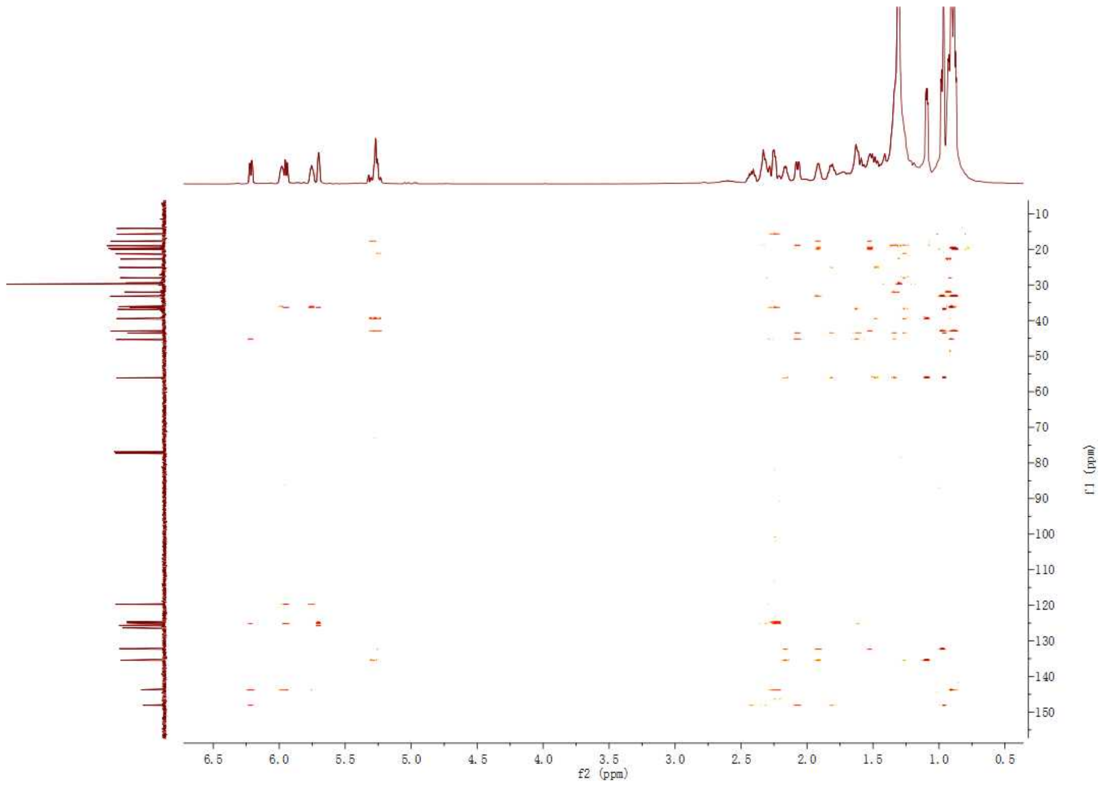


**Figure S26**. HMBC spectrum of ergosterene (**8**).

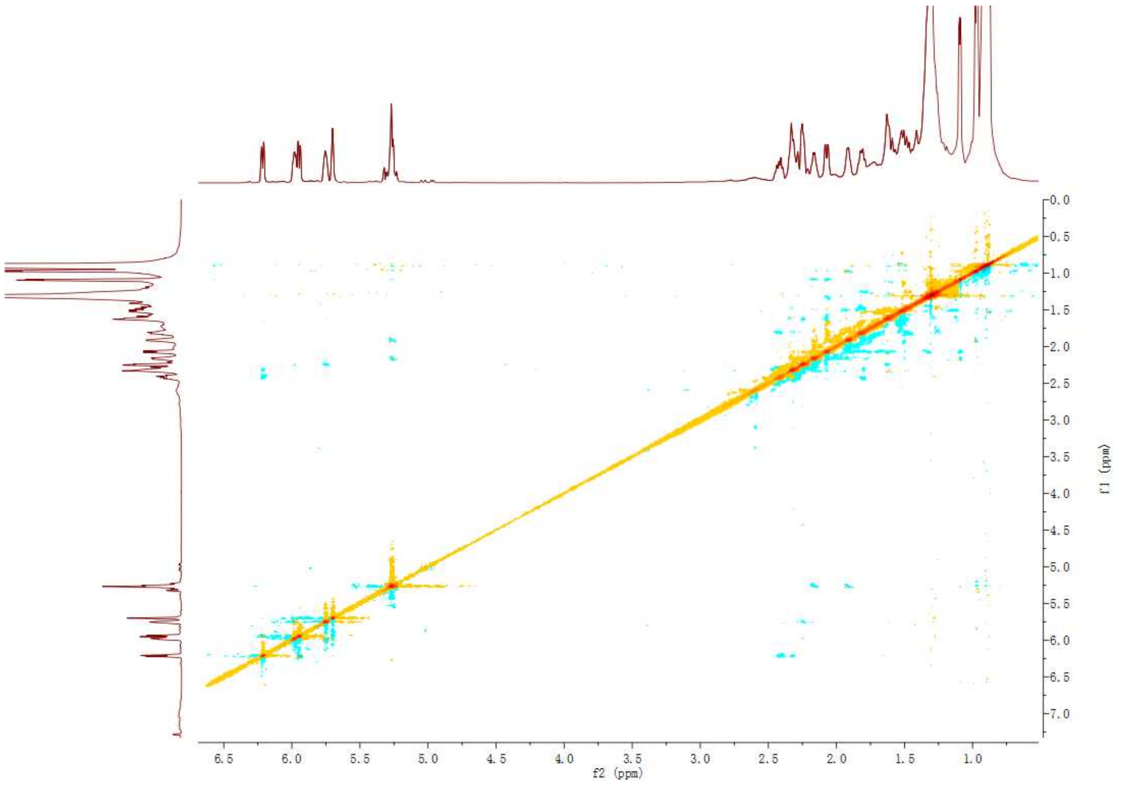


**Figure S27**. ROESY spectrum of ergosterene (**8**).

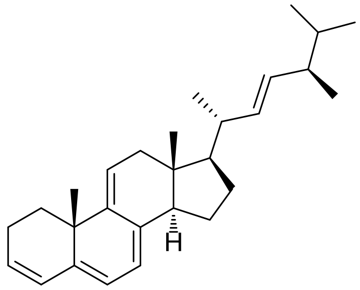

(22E)-ergosta-3,5,7,9(11),22-pentaene (8)

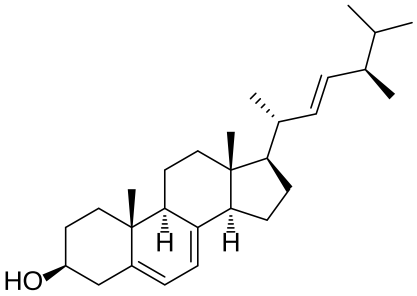

ergosterol

**Figure S28**. Structures of ergosterene (**8**) and ergosterol.
